# Spinal injury induces a stem cell-like progenitor state that promotes regenerative neurogenesis via *clcf1* in zebrafish

**DOI:** 10.64898/2026.06.25.734505

**Authors:** Markus Westphal, Rachel Branch, Alberto Docampo-Seara, Mehmet Ilyas Cosacak, Ekaterina Logvinova, Maia Bellia, Shaoyun Cheng, Themistoklis M. Tsarouchas, Anja Bretschneider, Daniela Zoeller, Ruoqi Shen, Luisa Scucces, Denise Bärhold, Thomas Becker, Catherina G. Becker

**Author notes:** equal contribution (Markus Westphal, Rachel Branch). senior authors (Thomas Becker, Catherina G. Becker).

## Abstract

After spinal injury, zebrafish, in contrast to mammals, show regenerative neurogenesis, characterized by enhanced injury-induced proliferation of ependymo-radial glial cells (ERGs) and an increase in injury-induced generation of neurons from these progenitors. It is unclear whether regenerative neurogenesis simply recapitulates development or uses regeneration-specific mechanisms. Using scRNA-seq and *in vivo* validation we find a spinal injury-induced state in ERGs (iiERGs) in larval zebrafish. This cell state emerges mostly without proliferation and has stem cell characteristics, including weak expression of neurogenic genes and strong expression of stemness factors, such as *lin28a*. Expression of *lin28a* is not detectable during ongoing developmental neurogenesis. Following spinal cord lesion, *lin28a* disruption increases the numbers of ERGs undergoing neuronal differentiation and of newly-generated neurons, at the expense of proliferating ERGs and iiERGs. This supports a stemness-preserving role of *lin28a* in iiERGs. Importantly, iiERGs secrete growth factors, including the regeneration-specific cytokine *clcf1,* which depends in part on *lin28a* expression. Disruption of *clcf1* signalling impairs spinal progenitor proliferation and injury-induced generation of new neurons, but does not affect the emergence of iiERGs. Over-expression of *clcf1* is sufficient to augment neurogenesis in unlesioned animals without inducing the iiERG state, indicating that *clcf1* acts as a generic growth factor. Hence, we describe an injury-specific stem cell-like ERG population that regulates regenerative neurogenesis by attenuating neuronal differentiation via *lin28a* and promoting progenitor proliferation via *clcf1*.

## INTRODUCTION

Spinal cord injury causes extensive and prolonged localized loss of neurons near the lesion (Casella et al., 2006; Dusart & Schwab, 1994). Lost neurons are never replaced (Meletis et al., 2008). In rodent species, although spinal progenitor cells display stem cell-like properties and proliferate after injury, they give rise to reactive astrocytes that contribute to the formation of an inhibitory glial scar (Barnabe-Heider et al., 2010; Johansson et al., 1999; Meletis et al., 2008; Sabelstrom et al., 2014; Silver & Miller, 2004; Weiss et al., 1996). Spinal progenitors can be experimentally reprogrammed, e.g. to produce more oligodendrocytes (Llorens-Bobadilla et al., 2020), but generation of neurons has not been reported.

In contrast, anamniote species such as zebrafish exhibit robust spinal cord regeneration (Becker et al., 2004; Becker et al., 1997; Mokalled et al., 2016). In larval and adult zebrafish, spinal ependymo-radial glial cells (ERGs) augment neurogenesis following a lesion and replace lost neurons (Becker & Becker, 2015; Briona et al., 2015; Mokalled et al., 2016; Ohnmacht et al., 2016; Reimer et al., 2008). After a spinal lesion, domain-specific ERGs proliferate and generate distinct neuronal subtypes (Kuscha et al., 2012a; Reimer et al., 2008). Ventromedial *olig2*-expressing ERGs from the motor neuron progenitor (pMN) domain, for example, produce *mnx1*-expressing motor neurons during regenerative neurogenesis, resembling development (Reimer et al., 2008). While in adult zebrafish, developmental neurogenesis has largely ceased, in larval zebrafish neurogenesis is ongoing, but an injury induces, for example, a ∼4-fold increase in motor neuron production from pMN ERGs (Ohnmacht et al., 2016). Hence in larval zebrafish, regenerative neurogenesis can directly be compared to ongoing developmental neurogenesis.

Spinal cord lesion reinitiates key developmental signaling pathways in pMN-domain ERGs, including Notch (Dias et al., 2012), Wnt (Briona et al., 2015), Fgf (Goldshmit et al., 2012), Egf (Cigliola et al., 2023) and Shh (Reimer et al., 2009) signaling, with dopamine (Reimer et al., 2013) and serotonin (Barreiro-Iglesias et al., 2015) signaling further reinforcing this regenerative response. Importantly, beyond these developmental cues, immune cells provide essential lesion-specific activation signals that may not recapitulate development. For example, macrophage-derived Tnfa induces AP-1 activity in ERGs, thereby promoting regenerative neurogenesis (Cavone et al., 2021) and microglia-derived Sema4ab attenuates regenerative neurogenesis (Docampo-Seara et al., 2026). Together these results suggest that ERGs after a spinal cord injury integrate both developmental and injury-specific signals. ERGs clearly engage in regenerative neurogenesis after spinal cord injury in zebrafish, but the injury-induced cellular states in ERGs that enable this response remain unknown. In particular, it remains unknown whether a specific stemness or activation state exists that permits a regenerative competence in ERGs. Defining the activation cues and cellular states that characterize regenerative ERGs may reveal strategies to restore neurogenic potential in endogenous spinal progenitors in mammals or to enhance the efficacy of transplanted neural stem cells for functional repair (Ceto et al., 2020). We therefore hypothesized that spinal cord injury triggers a unique ERG state that is molecularly and functionally different from developmental ERG states and is essential for enabling regenerative neurogenesis.

Here, we reveal a distinct transcriptional state in ERGs after lesion, characterized by the expression of both stemness-related markers and secreted factors not expressed in the unlesioned spinal cord, which nevertheless undergoes low levels of developmental neurogenesis. We name this population/state injury-induced ERGs (iiERGs). Disruption of the iiERG stemness factor *lin28a* results in increased regenerative neurogenesis, accompanied by a depletion of progenitors after spinal cord lesion. Disruption of the iiERG-expressed cytokine gene *clcf1,* as well as disruptions of its receptor *il6st*, reduced progenitor proliferation and decreased regenerative neurogenesis. These findings suggest a model in which spinal cord injury induces unique stemness characteristics in a subpopulation of ERGs, which in turn promotes regenerative neurogenesis through Clcf1/Il6st signaling.

## RESULTS

### A Spinal Cord Lesion Induces Delayed Neurogenesis

In larval zebrafish, a lesion of the spinal cord at 3 days post-fertilization (dpf) leads to regenerative neurogenesis that peaks at 48 hours post-lesion (hpl), coinciding with axon regrowth and recovery of swimming function (Ohnmacht et al., 2016). To analyse the temporal dynamics of lesion-induced neurogenesis in this system, we used *mnx1:GFP* transgenic animals, in which motor neurons are labeled with green fluorescent protein (GFP) (Flanagan-Steet et al., 2005). Motor neurons are the primary neuronal cell type generated in response to spinal cord lesion alongside interneurons (Kuscha et al., 2012a; Ohnmacht et al., 2016; Reimer et al., 2009; Reimer et al., 2008).

To determine when regenerative neurogenesis begins after injury, we performed a “birthdating” analysis of lesion-induced motor neuron generation. To this end, we lesioned animals at 3 dpf and pulse-labeled them with EdU at various post-lesion time points for 30 minutes each before determining the cumulative number of EdU-positive motor neurons near the lesion at 48 hpl (Figure 1A). Because only cells undergoing DNA synthesis at the time of the EdU pulse become labeled, this approach identifies the time points at which progenitors were proliferating prior to differentiation into motor neurons. We observed the first increase in EdU-labeled motor neurons after EdU administration at 16 hpl (3.6-fold), with a further increase after EdU-labeling at 24 hpl (6.9-fold), as compared to baseline developmental neurogenesis in age-matched unlesioned larvae (Figure 1A). These data showed that progenitors contributing to regenerative motor neurogenesis first entered the cell cycle at around 16 hpl suggesting a roughly 16-hour delay before the onset of regenerative neurogenesis.

**Fig. 1.**
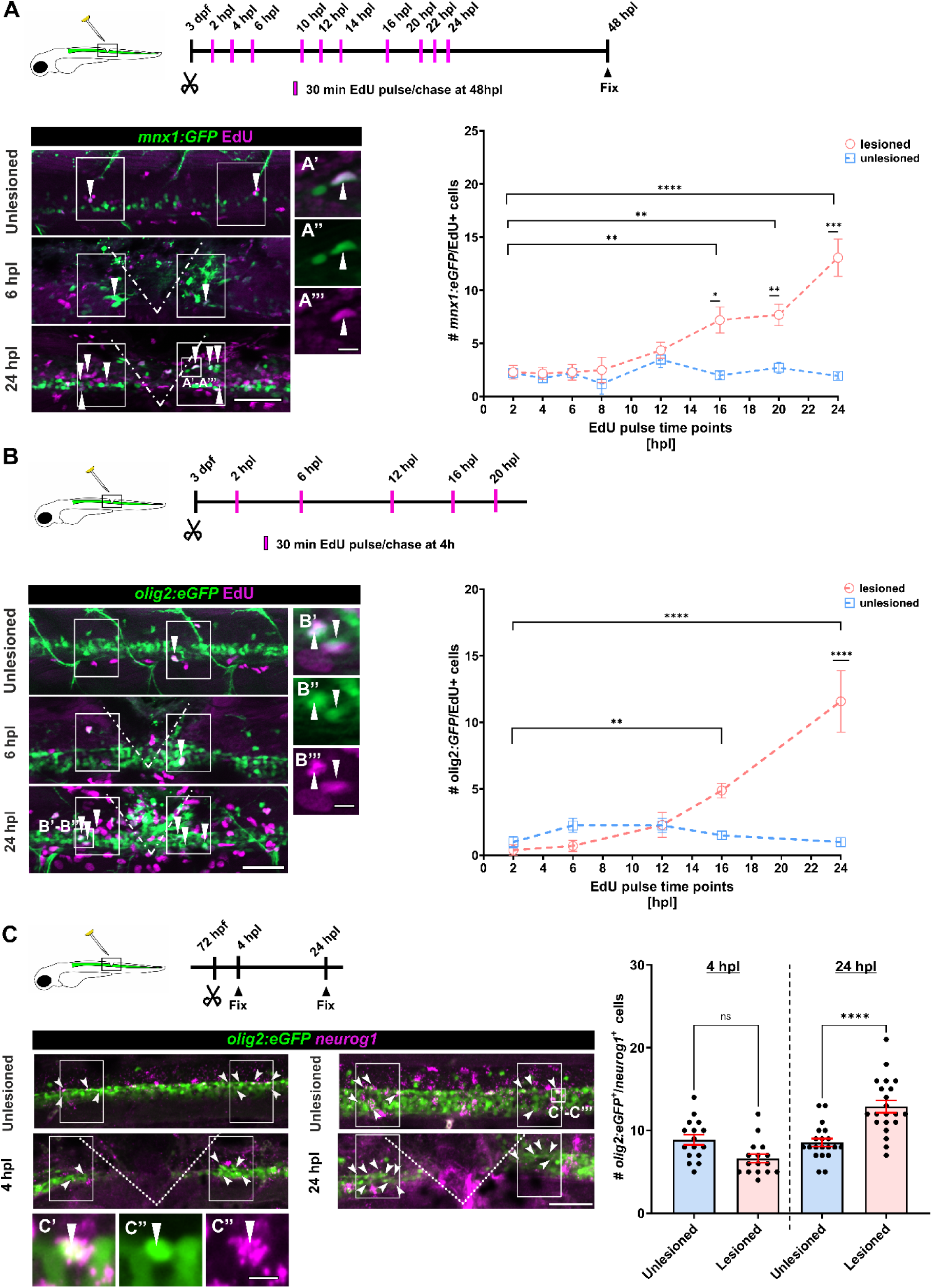
A spinal cord lesion induces delayed neurogenesis. **(A)** EdU pulse at different time points and chase at 48 hpl in *mnx1:GFP* transgenic animals shows a delay in the generation of new born neurons after injury. Arrows point at double positive cells (Two-way ANOVA: p < 0.0001; Holm-Sídák’s multiple comparaison test: p* = 00124, p** = 0.0030, p*** = 0.0001)**. (A’-A’’’)** High magnifications show an example of a *mnx1:GFP*/EdU double-positive cell. **(B)** EdU pulses at different time points and detection 4h later in *olig2:eGFP* animals shows a delay in the lesion-induced increase in the number of proliferating motor neuron precursors. Arrows point at double positive cells (Two-way ANOVA: p < 0.0001; Holm-Sídák’s multiple comparaison test: p** = 0.0073, p*** < 0.0001)**. (B’-B’’’)** High magnifications showing an example of an *olig2:eGFP*/EdU double positive cell. **(C)** The number *olig2:eGFP /neurog1* double-positive ERGs is increased at 24 hpl, but not at 4 hpl, compared to age matched unlesioned controls (One Way ANOVA: p < 0.0001; Tukey’s multiple comparison test: p**** < 0.0001)**. (C’-C’’’)** High magnifications showing an example of *olig2:eGFP/neurog1* double-positive cell. **Scale bars:** 50 µm (A, B, C), 10 µm (A’-A’’’, B’-B’’’, C’-C’’’). Dorsal is up and rostral is left.

To validate the proliferative dynamics inferred from the motor neuron birthdating experiment, we analysed acute proliferation patterns in an *olig2* reporter line (*olig2:eGFP*) that labels pMN cells in the ventral spinal cord, the source of motor neurons both during development and regeneration (Park et al., 2004; Reimer et al., 2008). To label only acutely proliferating progenitors, we applied EdU at similar time points as above to label new motor neurons, but quantified EdU incorporation in *olig2*^+^ cells shortly after pulse-labeling (4 h delay) rather than at later stages following neuronal labelling (Figure 1B). This experiment revealed increased EdU incorporation in *olig2:eGFP^+^* cells first at 16 hpl (3.3-fold) and a marked increase at 24 hpl (11.6-fold) in lesioned larvae, compared to age-matched unlesioned larvae (Figure 1B). These time points matched those identified with the birthdating analysis, indicating that progenitors giving rise to regenerated motor neurons first entered the cell cycle at these stages. Therefore, the temporal dynamic of acute proliferation of pMN progenitors corroborated our birthdating results and these findings indicate that progenitor proliferation in the lesioned spinal cord was increased from 16 hpl onwards.

Next, we assessed whether neuronal commitment of ERGs was changed after spinal cord lesion. To that end, we determined expression of *neurog1* in pMN progenitors. Neurogenins, such as the pro-neural bHLH transcription factor *neurog1,* are required for neurogenesis and for motor neuron precursor commitment in the ventral spinal cord during development (Bertrand et al., 2002; Lee et al., 2005; Scardigli et al., 2001; Velasco et al., 2017). To assess whether *olig2:eGFP^+^* progenitors in the lesioned spinal cord exhibited an increased commitment to neurogenesis, we quantified the number of *neurog1*-expressing *olig2:eGFP^+^* pMN progenitors before lesion-induced neurogenesis at 4 hpl and during lesion-induced neurogenesis at 24 (Figure 1C). We found that the number of *neurog1^+^*/*olig2:eGFP^+^* cells was not elevated at 4 hpl, but was increased at 24 hpl (1.5-fold) in lesioned larvae, compared to age-matched unlesioned larvae (Figure 1C). This delayed increase coincided with the lesion-induced generation of motor neurons. Together, these observations demonstrated that after a spinal cord lesion, a robust increase in progenitor proliferation, neuronal commitment, and neurogenesis occur with roughly a 16-hour delay after lesion.

### scRNA-seq Identifies an Injury-Specific Cluster of ERGs

To identify lesion-induced changes in ERGs that may occur during the initial delay phase, we performed single-cell RNA sequencing (scRNA-seq) on FACS-sorted *her4.1:eGFP^+^* cells, which predominantly labeled ERGs (Dias et al., 2012; Yeo et al., 2007) from lesioned larvae at 12 hpl and age-matched unlesioned controls. FACS-sorting enriches the samples for *her4.1:eGFP^+^* ERGs, but also their neuronal subtype-committed progeny that retain eGFP fluorescence after being generated from GFP-expressing ERGs (Cavone et al., 2021). We selected 12 hpl as the time point just prior to the onset of regenerative neurogenesis to most likely capture early and late lesion-induced ERG states.

For scRNAseq profiles, we pooled cells from 200 animals each, and obtained 7598 cells for the 12 hpl lesioned sample, and 7264 cells for the unlesioned age-matched sample, after quality control. Unsupervised clustering identified 36 clusters that fell into 15 main categories of neural cell types (Figure 2A; Supplementary Figure 1). We identified one cluster as ERGs by highest expression levels of the ERG-specific genes *her4.1 and foxj1a* (Ribeiro et al., 2017; Yeo et al., 2007), which define distinct ERG subpopulations through their mostly non-overlapping expression patterns, and together capture ERG heterogeneity of the spinal cord (Dias et al., 2012; Ribeiro et al., 2017) (Figure 2B, C). Sub-clustering of ERGs yielded 943 cells for the 12 hpl lesioned sample, and 605 cells for the unlesioned age-matched sample, and identified 11 ERG clusters that represented 9 main cell types (Figure 2D). These clusters represented a heterogenous population of dorsal progenitor ERGs (dERGs) enriched in marker *pax7a, zic1, zic2a, olig3, msx1a* (Briscoe et al., 2000; Rondon et al., 2025), p1 progenitor ERGs (p1ERGs), enriched in marker *irx3a and dbx1a* (Briscoe et al., 2000), lateral floor plate (lfpERGs), enriched in marker *shha* and *nkx2.9* (Charrier et al., 2002; Park et al., 2004), pMN progenitor ERGs (pmnERGs) enriched in maker *olig2* and *olig1* (Lu et al., 2002; Park et al., 2004), actively proliferating ERGs (apERGs). enriched in markers *mki67* and *pcna* (Moldovan et al., 2007; Scholzen & Gerdes, 2000), neural progenitor ERGs (nERGs) enriched in marker *neurog1, neurod1 and notch1 (Bertrand et al., 2002; Katreddi et al., 2022)*, motor neuron committed ERGs (mnERGs) enriched in marker *mnx1and chata* (Arber et al., 1999) and an unidentified ERG cluster (NN) that does not share any domain specific markers enriched in marker *s100b1a1*, which is associated with ERG identity (Jurisch-Yaksi et al., 2020). Importantly, we observed a large cluster of ERGs that was almost exclusively present in the lesioned sample (0.5 % of all ERGs in the unlesioned sample vs. 30.3 % in the lesioned sample; Figure 2E, F), which we therefore termed injury-induced ERGs (iiERGs).

**Fig. 2.**
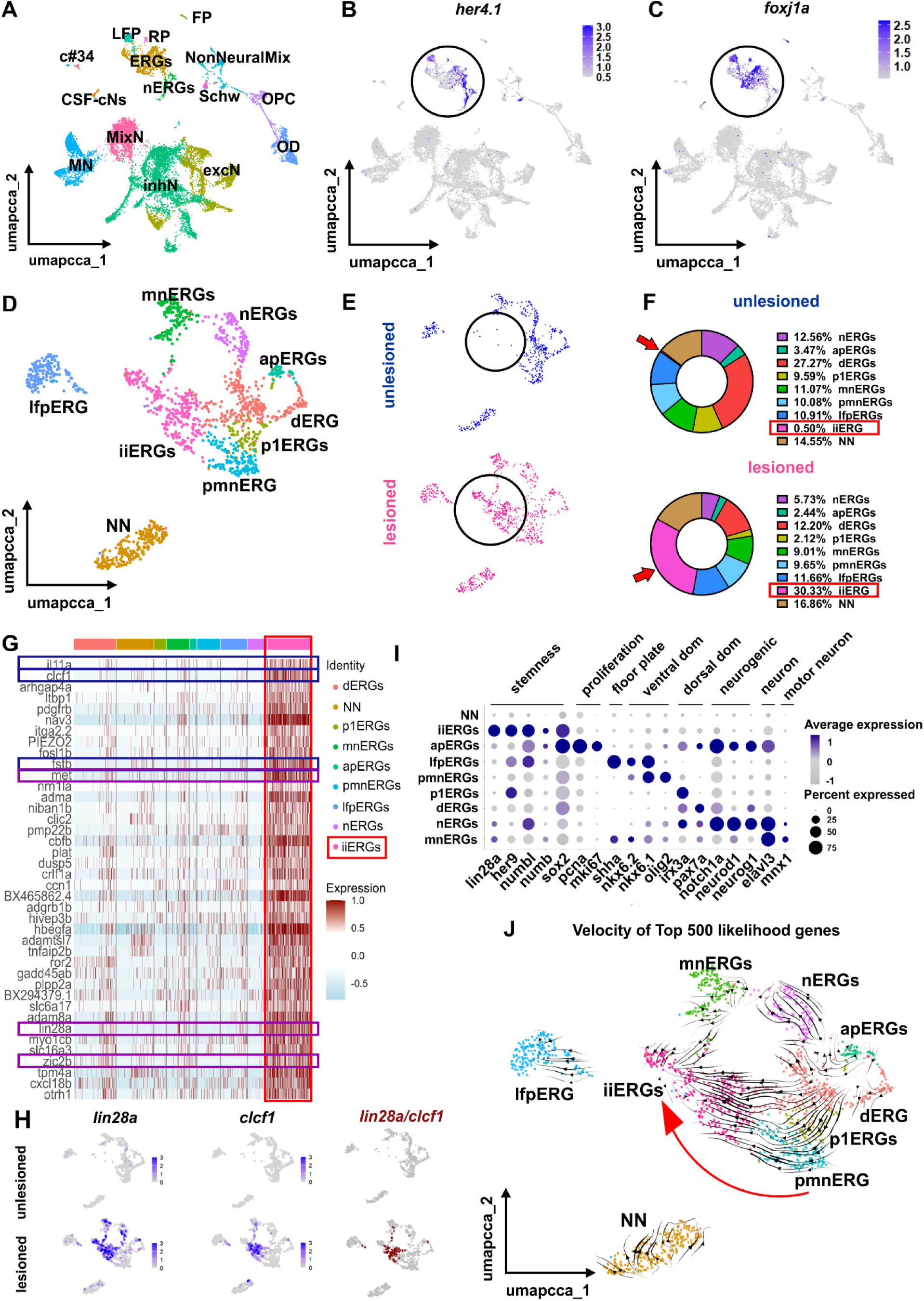
scRNAseq identifies an injury-induced ERG population. **(A)** Umap showing the main cell types in the lesioned spinal cord at 12 hpl. **(B, C)** Feature plots showing the expression pattern of ERG genes *her4.1* and *foxj1a*. Circle points the co-expressed clusters used for sub-setting. **(D)** Unsupervised sub-clustering revealed different clusters that fall into 9 main categories of cell types. **(E)** UMAPs comparing unlesioned and lesioned conditions showing that the iiERGs are almost non-existent before injury (circle). **(F)** scRNAseq quantification comparing unlesioned and lesioned samples, in which iiERGs (red arrows) represent a 0.5% of the total cells in unlesioned larvae and 30.33% after lesion (red box). **(G)** Top 40 ranked gene analysis for iiERG shows an enrichment of signalling factors such as *clcf1* (and cofactor *crlf1a*) and stemness factors such as *lin28a*. Blue boxes indicate secreted factors and purple boxes highlight stemness factors. **(H)** Feature plots showing that *lin28a* and *clcf1* expression is enriched in iiERGs, only present after injury, and co-localized in the same cells. **(I)** Dot plot showing marker genes for cell cluster annotation. Note that iiERGs do not show neurogenic, proliferation or domain specific markers, but the general progenitor marker *sox2* and stemness markers such as *numb, numbl*, and *her9*. **(J)** Velocity analysis of 500 likelihood genes showing trajectories from all ERG domain clusters to the iiERG cluster (red arrow).

An analysis of the most highly enriched transcripts in iiERGs (in the top 40 list of differential gene expression) indicated genes controlling stemness and self-renewal in embryonic stem cells, such as *lin28a* (Bin et al., 2012; Hanna et al., 2009; Yu et al., 2007), *fosl1b* (Pecce et al., 2021), *met* (Li et al., 2011), *zic2b* (Luo et al., 2015), and in neural stem cells, such as *her9* (Engel-Pizcueta et al., 2025)*, numb*, *numbl* (Knoblich et al., 1995) and *sox2* (Ellis et al., 2004). Highly enriched transcripts were also those related to secreted cytokines and other signaling factors, such as *il11a*, *clcf1* (Rose-John, 2018), and *fstb* (de Winter et al., 1996) (Figure 2G, H; Supplementary Figure 2A). A feature plot analysis revealed enriched expression of the stemness gene *lin28a* and the cytokine *clcf1* in iiERGs after the lesion and furthermore that *lin28a* and *clcf1* were co-expressed in iiERGs after the lesion (Figure 2H). Notably, some markers associated with pluripotency in Müller glia after retina injury, including *sox2* (Gorsuch et al., 2017)*, mycb* (Lee et al., 2024; Mitra et al., 2019)*, klf4* (Ramachandran et al., 2010)*, stat3, and lin28a* (Ramachandran et al., 2010) were strongly enriched in iiERGs, but other canonical pluripotency markers such as *pou5f3* or *nanog* (Ramachandran et al., 2010; Sharma et al., 2019) were not expressed in iiERGs (Supplementary Figure 2B), indicating clear differences between activated Müller cells and iiERGs. Additionally, neural stemness factors such as *her9 (Engel-Pizcueta et al., 2025), numb* and *numbl (Knoblich et al., 1995)*, were enriched in iiERGs, while markers for specific progenitor domains (such as dorsal progenitors, p1 and pMN domains) were expressed at very low levels (Figure 2I). Likewise, proliferation markers, such as *mki67*, *pcna,* as well as neuronal differentiation markers, such as *neurog1*, *neurod1 (Bertrand et al., 2002),* were hardly expressed in iiERGs (Figure 2I and Supplementary Figure 2C). A Velocity analysis of developmental trajectories based on the 500 genes with the highest likelihood scores, representing genes which splicing and transcriptional dynamics that best fit the RNA velocity model (see Material and Methods for details) (Figure 2J) indicated that neuronally differentiating ERGs were possibly derived from domain-specific ERGs via a proliferative state, as previously demonstrated for the adult zebrafish spinal cord *in vivo* (Kuscha et al., 2012b; Reimer et al., 2008). Another trajectory led from domain-specific ERGs to iiERGs without an intermediate proliferative state, suggesting potential de-differentiation of domain-specific ERGs to the iiERG state (Figure 2J, arrow). The analysis did not provide indications for a trajectory from iiERGs to neuronally differentiating ERGs. However, this might be due to the early time point (12hpl) analyzed, since our neuron birthdating analysis indicate that regenerative neurogenesis occurs after 16 hpl. These findings suggested that iiERGs initially arise from pre-existing domain-specific ERG populations. Together this data suggested that iiERGs represented an injury-specific neural stem cell-like state with a function in growth factor secretion.

### iiERGs are induced by spinal cord lesion *in vivo*

To characterize iiERGs *in vivo*, we first analysed the time-course of iiERG presence in the lesion vicinity. We used HCR-FISH for the specific marker *lin28a* in *her4.1:GFP* transgenic larvae, in which all ERGs are labelled by GFP (Yeo et al., 2007). Expression of *lin28a* was below detection level in the entire spinal cord of unlesioned controls, confirming the findings from our scRNA-seq data (Figure 3A, B). After lesion, *lin28a* was detectably expressed only in *her4.1:eGFP^+^*cells. At 2 hpl, the first *lin28a^+^*/*her4.1:eGFP^+^*cells were detected. We observed a peak in the number of *lin28a^+^*/*her4.1:eGFP*^+^ cells at 12 hpl, followed by a gradual decline in their number at 16 hpl, 24 hpl, 30 hpl, and 48 hpl (Figure 3 B, C). Transcript abundance was always strongest next to the lesion and extended symmetrically maximally ∼160 µm rostrally and caudally (measured at 12 hpl, when *lin28a*^+^/*her4.1:eGFP*^+^ cells peaked; Figure 3B, C). Thus, *lin28a* expression was restricted to ERGs in the vicinity of a spinal lesion, with a peak in expression at 12 hours post-injury, slightly before the beginning of regenerative neurogenesis at 16 hpl.

**Fig. 3.**
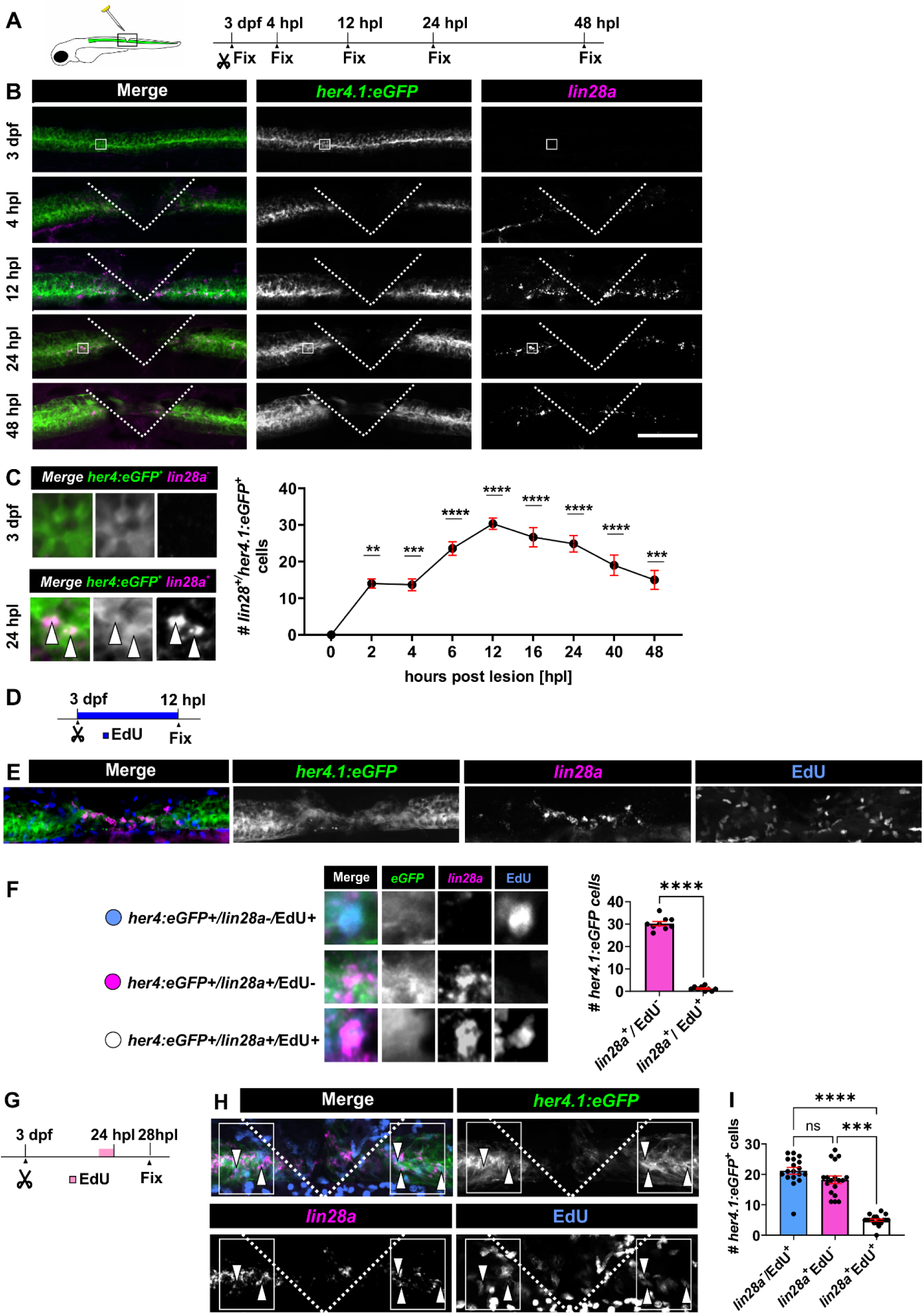
Lin28a is enriched in iiERGs, which show low proliferation. **(A)** Schematic of the experiment shown in B and quantified in C. **(B)** HCR-FISH against *lin28a* in *her4.1:eGFP* animals shows an upregulation of *lin28a* in ERGs after injury that decays over time. **(C)** Quantification shows that *lin28a* expression peaks around 12 hpl and decays thereafter (One-Way ANOVA: p < 0.0001; Tukey’s multiple comparison test: p*** < 0.001, p**** < 0.0001)**. (D)** Schematic of the experiment performed in E and quantified in F. **(E)** Example images of *her4.1:eGFP / lin28a /* EdU triple-labelling is shown. **(F)** Quantification shows that the vast majority of *lin28a^+^* cells are EdU-negative (*lin28a+/*EdU-: 30.2 ± 0.9 cells; *lin28a+/*EdU+: 1.2 ± 0.3 cells; Unparied T-test: p**** < 0.0001)**. (G)** Schematic of the experiment to measure acute proliferation in *lin28a*+ ERGs preformed in H. **(H)** Acutely proliferating ERGs are shown in *her4.1:eGFP / lin28a /* EdU triple-labelling. **(I)** Quantification shows that the vast majority of *lin28a+* cells are EdU negative (*lin28a^-^/*EdU^+^: 21.2 ± 1.1 cells; *lin28a^+^/*EdU^-^: 18.2 ± 1.2 cells; *lin28a^+^/*EdU^+^: 4.9 ± 0.4 cells; One-Way ANOVA: p < 0.0001; Tukey’s multiple comparison test: p*** = 0.0001, p**** < 0.0001)**. Scale bars:** 100 µm (B), 50 µm (E, H). Dorsal is up and rostral is left.

To test our computational trajectory analysis that domain-specific progenitors contribute to the iiERG pool after a spinal cord lesion, we performed *lin28a* HCR-FISH on *olig2:eGFP* larvae in which pMN progenitors are genetically labeled and the *olig2*-driven GFP signal can serve as a short-term lineage tracer due to persistent GFP stability and has been used as a lineage tracer in adult zebrafish spinal cords (Reimer et al., 2008). We performed the experiment at 12 hpl, when iiERG numbers peaked (Supplementary Figure 3A). We quantified the proportion of *lin28a^+^* iiERGs among the *olig2:eGFP^+^*pMN progenitors and found that 33.4% of *olig2:eGFP^+^* cells expressed *lin28a* (Supplementary Figure 3B). Our finding indicates that iiERGs arise, at least in part, from domain-specific pMN progenitors, which is consistent with the trajectories predicted by RNA Velocity analysis of the scRNA-seq data.

To determine whether iiERGs arose through proliferation of ERGs, we incubated larvae continuously with EdU from the lesion time point at 3 dpf to the peak of *lin28a* expression at 12 hpl and determined the proportion of cumulatively labelled EdU^+^ iiERGs at that timepoint (Figure 3D, E). We found that only ∼3 % of the iiERGs (*lin28a^+^/ her4.1:eGFP^+^*) had incorporated EdU during that time (Figure 3E, F), indicating that iiERGs mostly arose by de-differentiation of pre-existing ERGs, which was consistent with the RNA Velocity analysis of the scRNA-seq.

Next, we aimed to determine whether iiERGs were proliferating during motor neuron generation. We performed a single EdU pulse for 30 min at 24 hpl, a time point when proliferation and neurogenesis in the lesioned spinal cord peaked, and analyzed incorporation of EdU and *lin28a* expression in *her4.1:eGFP^+^*progenitors at 4 hours post-pulse to measure acute proliferative activity (Figure 3G). The majority of *lin28a^+^* ERGs (79%) were not incorporating EdU. Conversely, of the EdU+ proliferating ERGs, only 21% were *lin28^+^*, indicating low proliferative activity of iiERGs during the peak of neurogenesis (Figure 3H, I).

To determine whether *lin28a*^+^ iiERGs were neurogenic *in vivo*, we co-labeled *lin28a* with the neurogenic marker *neurog1* in *her4.1:eGFP^+^* progenitors at 24 hpl (Supplementary Figure 4A). In agreement with the data from our scRNA-seq data set, we found that most *lin28a*^+^/*her4.1:eGFP^+^* cells (79 %) did not co-express *neurog1*, and, conversely, most *neurog1*^+^/*her4.1:eGFP^+^* cells (72.3 %) were *lin28a^-^* (Supplementary Figure 4B, C). Of note, the proportion of *lin28a*^+^/*her4.1:eGFP^+^* that expressed *neurog1* (21 %) was similar to that of EdU incorporation in these cells and may be evidence for low-level neuronal divisions in iiERGs. Overall, this suggested a relatively low neurogenic commitment in *lin28a^+^* ERGs, in line with a stemness-preserving function of *lin28a*.

### *lin28a* expression preserves the ERGs

To demonstrate potential stem cell features of iiERGs we decided to disrupt *lin28a*, because presence of *lin28a* is known to protect the neural progenitor pool, e.g. in human embryonic stem cell cultures and in the developing mouse cortex (Cimadamore et al., 2013; Yang et al., 2015). At the same time, its expression is highly enriched in iiERGs and no other parenchymal cells. Therefore, any effects of experimental gene disruption could be mostly attributed to iiERGs.

To abrogate *lin28a* function, we generated somatic CRISPR mutants (crispants) in which highly efficient gene disruption is achieved by injection of synthetic gRNAs targeting the gene together with Cas9 protein into 1-cell stage embryos (Docampo-Seara et al., 2026; Keatinge et al., 2021; Kroll et al., 2021). For somatic mutation, we used two specific gRNAs targeting exon 2, which encode the RNA-binding Cold Shock domain (Moss et al., 1997; Moss & Tang, 2003). Efficient disruption of the targeted site in injected larvae was confirmed by restriction fragment length polymorophism (RFLP) analysis, which indicated that the targeted restriction enzyme recognition sequence became resistant to digestion in DNA isolated from 3 dpf larvae (Supplementary Figure 5A). Furthermore, because *lin28a* was not expressed in unlesioned larvae, we performed qPCR analysis of *lin28a* expression in dissected trunks from spinal cord-lesioned and *lin28a* gRNA-injected somatic mutants and found a 55 % reduction in the abundance of *lin28a*-related transcripts compared to control-injected larvae (Supplementary Figure 5B). As an independent control, we used gRNAs with sequences that were not related to those used above, targeting *lin28a* exon 3 (Supplementary Figure 5A). All results were compared to those in larvae injected with unspecific scrambled gRNAs (Keatinge et al., 2021). At 5 days post-lesion (dpl), somatic *lin28a* mutants developed with normal body shape, though they exhibited a slight reduction in body length (7.5 %) and eye diameter (5.3 %) compared to control gRNA-injected larvae (Supplementary Figure 5C-F).

To assess the impact of *lin28a* disruption on ERG maintenance and proliferation, we performed spinal lesions at 3 dpf, followed by cumulative EdU labeling for 48 hours using the *her4.1:GFP* transgenic line (Figure 4A). In unlesioned animals, *lin28a* disruption did not affect EdU incorporation in *her4.1:eGFP^+^* cells (Figure 4B), which was consistent with the absence of *lin28a* expression during these larval stages. This also serves as a control for potential non-specific effects of the gRNAs on neurogenesis. After lesion, control larvae exhibited a 31 % increase in the number of EdU+ ERGs compared to unlesioned control animals (Figure 4B), consistent with lesion-induced neurogenesis, as previously reported (Ohmnacht et al., 2016). Lesioned *lin28a* somatic mutants showed a 52 % reduction in the number of EdU^+^ ERGs, compared to lesioned control larvae (Figure 4B). Single-pulse EdU treatment (30 min at 24 hpl, analysis at 28 hpl), also indicated a 36 % reduction in acute proliferation of these cells in lesioned somatic mutants for *lin28a* (Figure 4C, D). Hence, *lin28a* disruption reduced the number of proliferative progenitor cells either by reducing progenitor proliferation and/or increasing neuronal differentiation.

**Fig. 4.**
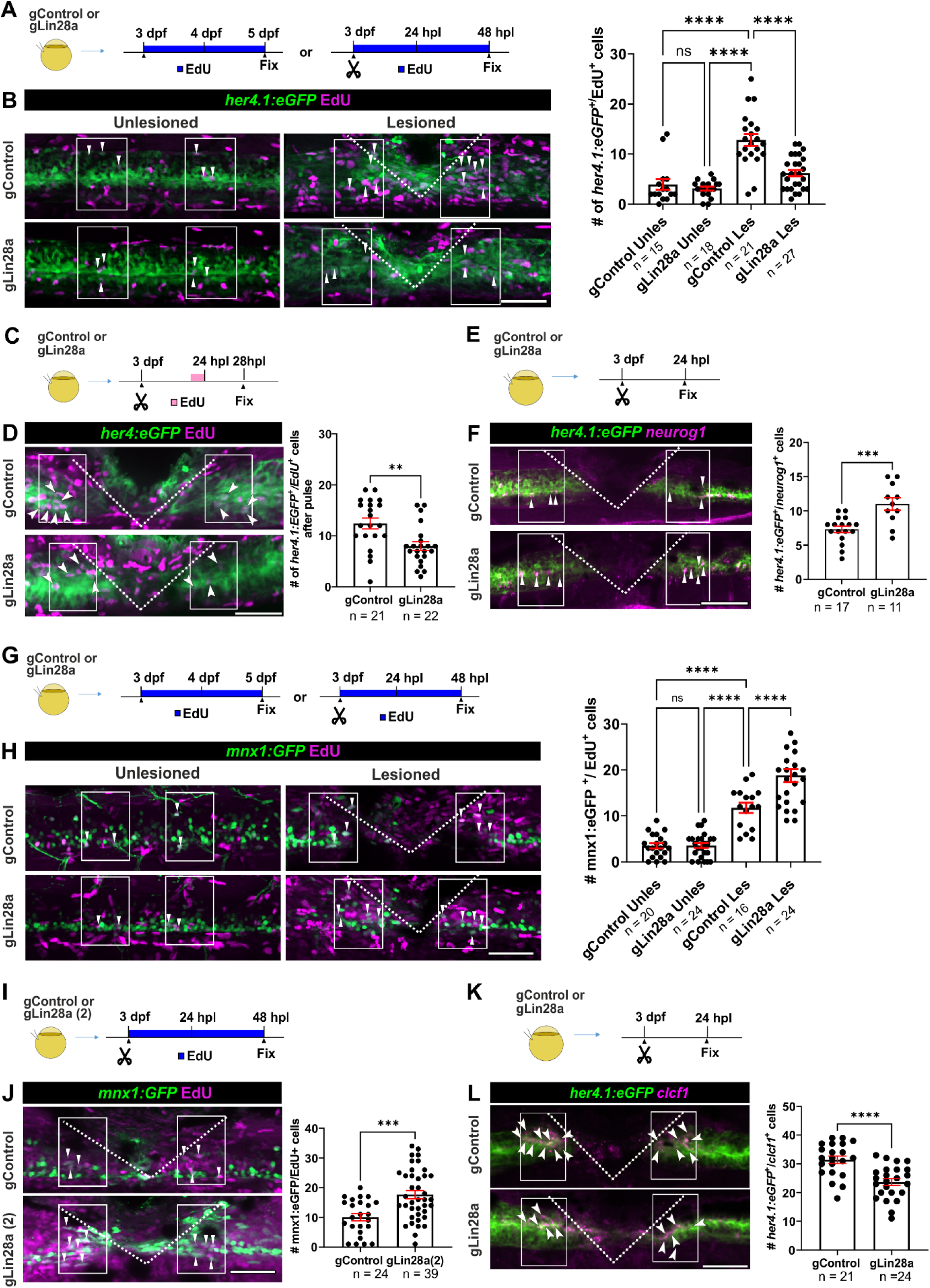
*lin28a* loss-of-function increases regenerative neurogenesis by depleting the progenitor pool. **(A-B)** (A) shows schematic of the experiment. (B) Ongoing developmental ERG proliferation (*her4.1:eGFP*^+^/EdU^+^) is unchanged in unlesioned somatic mutants for *lin28a*, but lesion-induced generation of ERGs is reduced in somatic mutants for *lin28a* compared to control gRNA-injected larvae (gControl Unles: 3.9 cells per larva ± 1.1; gLin28a Unles: 3.1 cells per larva ± 0.4; gControl Les: 12.8 cells per larva ± 1.2; gLin28a Les: 6.2 cells per larva ± 0.6; Two-way ANOVA: p < 0.0001; Holm-Šidák’s multiple comparison test: p < 0.0001). **(C-D)** (C) shows a schematic of the experiment. (D) Lesion-induced ERG activation (*her4.1:eGFP^+^/EdU^+^, pulse)* is decreased in lesioned somatic mutants for *lin28a* (gControl: 12.4 cells per larva ± 1.1; gLin28a: 8.0 cells per larva ± 0.8; Mann-Whitney U-test: p = 0.0020). **(E-F)** (E) shows a schematic of the experiment. (F) Lesion-induced neurogenic progenitors (*her4.1:eGFP^+^/neurog1^+^)* are increased in number in somatic mutants for *lin28a* at 24 hpl (gControl: 7.3 cells per larva ± 0.4; gLin28a: 11.0 cells per larva ± 0.9; Unpaired t-test: p = 0.0005). **(G-H)** (G) shows a schematic of the experiment. (H) Ongoing generation of motor neurons (*mnx1:GFP*^+^/EdU^+^) is unchanged in unlesioned somatic mutants for *lin28a*, but lesion-induced generation of motor neurons is increased in somatic mutants for *lin28a* (gControl Unles: 3.5 cells per larva ± 0.5; gLin28a Unles: 0.5 cells per larva ± 2.6; gControl Les: 11.75 cells per larva ± 1.1; gLin28a Les: 18.8 cells per larva ± 1.4; Two-way ANOVA: p < 0.0001; Holm-Šidák’s multiple comparison test: p < 0.0001). **(I-J)** (I) shows schematic of the experiment. (J) Lesion-induced generation of motor neurons (*mnx1:GFP*^+^/EdU^+^) is increased in somatic mutants for *lin28a* using alternative gRNAs to those used in G-H (gControl: 10.1 cells per larva ± 1.2; gLin28a: 17.7 cells per larva ± 1.3; Unpaired t-test: p = 0.0003). **(K-L)** (K) shows a schematic of the experiment. (L) HCR-FISH against *clcf1* in *her4.1:eGFP* shows a reduction in *lin28a* gRNA-injected larvae (gControl: 31.4 cells per larva ± 1.3; gLin28a: 23.7 cells per larva ± 1.2; Unpaired t-test: p < 0.0001). **Scale bars:** 50 µm. Dorsal is up and rostral is left.

To determine potential effects of *lin28a* disruption on progenitor differentiation we quantified ERGs that expressed the neuronal differentiation factor *neurog1* (Figure 4E). We observed a 50 % increase in the number of committed neurogenic precursors ERGs (*neurog1^+^/her4.1:eGFP^+^*) in *lin28a* somatic mutants, compared to lesioned control animals at 24 hpl (Figure 4F). This indicated that neuronal differentiation of ERGs was likely enhanced in the absence of *lin28a* function.

To investigate whether enhanced neuronal differentiation would also lead to increased production of new neurons during regeneration in *lin28a* somatic mutants, we analyzed motor neuron generation by cumulative EdU labeling in a transgenic *mnx1:GFP* motor neuron reporter line at 48 hours post-lesion, compared to age-matched unlesioned controls (Figure 4G). In unlesioned larvae, disrupting *lin28a* had no effect on the number of EdU^+^/*mnx1:GFP*^+^ motor neurons during ongoing developmental neurogenesis (Figure 4H). After spinal cord lesion, control gRNA-injected larvae showed a 3-4-fold increase in newly generated motor neurons (EdU^+^/*mnx1:GFP*^+^); Figure 4H), as reported previously (Ohmnacht et al., 2016). In lesioned *lin28a* somatic mutants, a further 60 % (gRNA combination 1; Figure 4H) increase in the number of newly generated motor neurons was observed. To corroborate this effect, we independently tested a second gRNA combination targeting *lin28a* gene in lesioned larvae, which similarly resulted in a 75% increase in the number of newly generated motor neurons after spinal cord (Figure 4I, J). These findings indicated that *lin28a* acted as a negative regulator of neuronal differentiation and thus likely preserved the ERG progenitor pool.

Loss of *lin28a* could also disrupt the iiERG state, similar to its necessity to induce a regenerative state in Müller glia in the injured retina (Ramachandran et al., 2010). This could prevent expression of relevant pro-proliferative factors for neural progenitor cells, such as *clcf1* (Zhao et al., 2014), which is highly enriched in expression in iiERGs, together with its obligatory co-factor *crlf1a* (Elson et al., 2000; Perret et al., 2004) in our scRNA-seq results. Therefore, we examined the expression of *clcf1* in *lin28a* somatic mutants (Figure 4K). At 24 hpl, we observed *clcf1* expression in *her4.1:eGFP^+^*ERGs in lesioned control larvae. This signal was reduced by 25% in *lin28a* somatic mutant compared to lesioned controls, suggesting reduced acquisition of the iiERG state (Figure 4L). Overall, reduced EdU labelling of ERGs, increased *neurog1* expression, and increased EdU labelling of motor neurons after gene disruption was consistent with a role of *lin28a* in preserving the progenitor pool by attenuating neuronal differentiation of iiERGs.

### iiERGs dynamically express *clcf1 in vivo*

To determine whether *clcf1* from iiERGs could indeed promote progenitor proliferation, we first mapped *in vivo* expression of the gene in detail (Figure 5A, B). No *clcf1* transcripts were detectable in the unlesioned spinal cord at 3 dpf. Following injury, *clcf1* expression in ERGs was first detected at 2 hpl, increasing progressively from 6 hpl to peak at 12 hpl. Thereafter, a continuous decline of the signal was observed up to the last point of analysis at 48 hpl (Figure 5B, C). The signal was centered around the lesion site and extended maximally ∼190 µm rostral and caudal to the lesion (measured at 12 hpl, when *clcf1*^+^/*her4.1:eGFP*^+^ cells peaked; Figure 5B, C). The time course and spatial pattern of *clcf1* expression resembled that of *lin28a* expression and the genes were co-expressed in ventricular cells (Supplementary Figure 6A), confirming results from scRNA-seq (cf. Figure 3B, C). Notably, there was also *clcf1* expression in the non-neural lesion site. This expression occurred mainly in fibroblasts as indicated by labelling of *clcf1* in the *pdgfrb:GFP* fibroblast reporter (Supplementary Figure 6B).

**Fig. 5.**
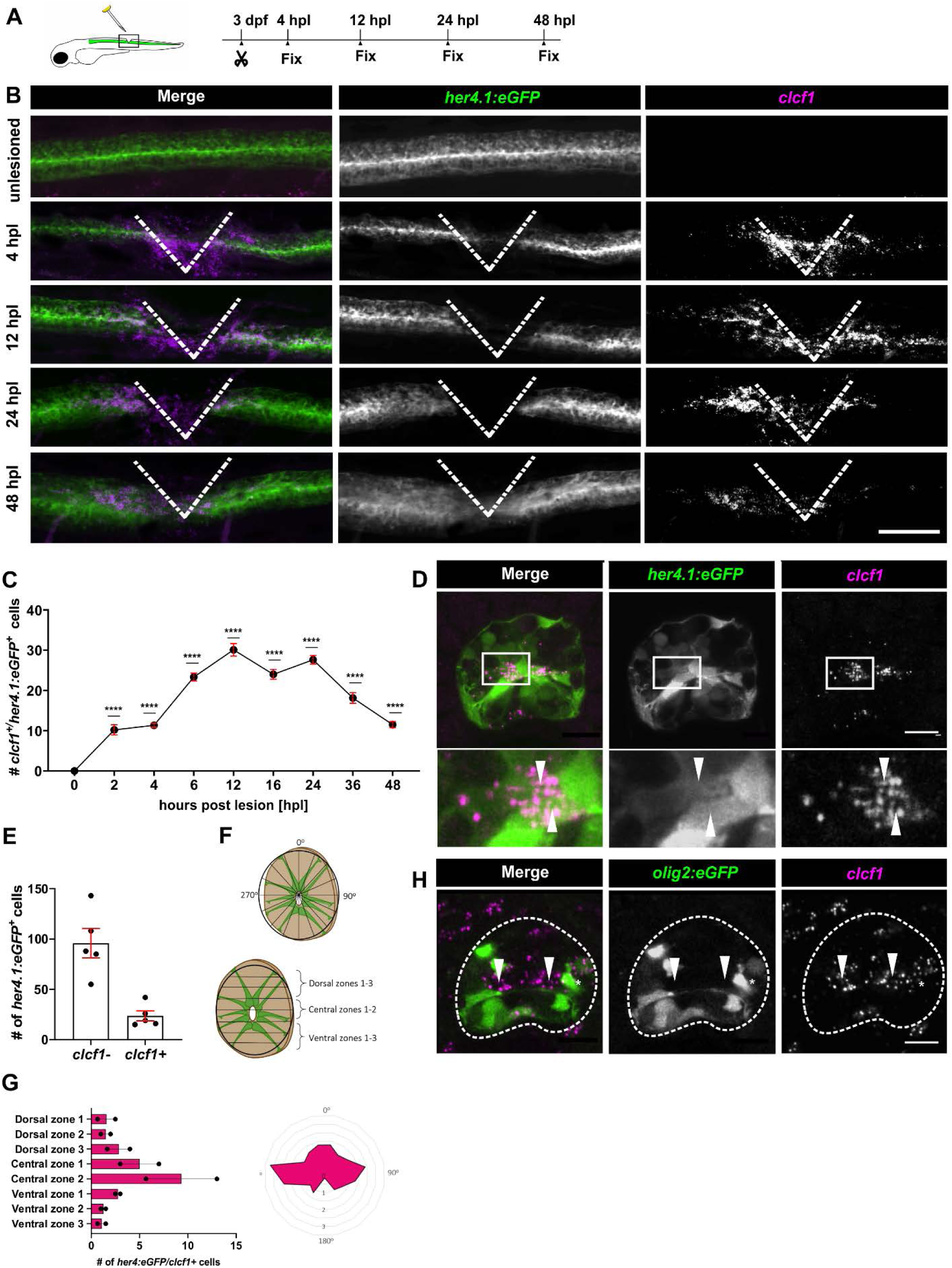
*clcf1* expression is enriched in iiERGs. **(A)** Schematic of the experiment shown in B. **(B, C)** HCR-FISH and quantification of *clcf1* in *her4.1:eGFP* animals shows the time course of *clcf1* expression. **(D)** Cross section of an HCR-FISH against *clcf1* in *her4.1:eGFP* larvae show colocalization in eGFP^+^ cells located in the centre of the spinal cord. **(E)** Quantification showing that *clcf1^+^/her4.1:eGFP^+^* cells represent a fraction of the total ERGs. **(F, G)** Spatial analysis of *clcf1^+^/her4.1:eGFP^+^* shows a preferential distribution in central zones around the central canal. **(H)** Cross section of a spinal cord processed for HCR-FISH against *clcf1* in an *olig2:eGFP* animal shows colocalization around the central canal and strong signal dorsal to the *olig2:GFP* cells**. Scale bars:** 100 µm (B), 15 µm (D, H). Dorsal is up and rostral is left.

Analysis of *clcf1* expression in cross sections of the spinal cord in *her4.1:eGFP* transgenic animals at 24 hpl showed that there was hardly any *clcf1* signal outside the ventricular zone and that 22 % of all *her4.1:eGFP^+^* ERGs were also *clcf1^+^* (Figure 5D, E). The highest number of *clcf1^+^/her4.1:eGFP^+^*cells were located at a median level of the spinal cord ((Figure 5F, G). Labelling of *clcf1* in the *olig2:GFP* reporter line indicated double labeled *clcf1*^+^/*olig2:GFP^+^*cells and a strong *clcf1* signal just dorsal to the *olig2:GFP^+^*pMN motor neuron progenitor domain at 24 hpl (Figure 5H). Hence, we detected lesion-induced presence of *clcf1^+^* iiERGs with highest numbers adjacent to the pMN domain, which gives rise to the motor neurons. iiERGs were thus ideally positioned for influencing the activity of pMNs by diffusible factors such as Clcf1 during regeneration.

### Clcf1 Promotes Regenerative Neurogenesis

To investigate the potential function of Clcf1 signaling in neurogenesis, we generated somatic and stable germline mutants. For somatic mutants, we used highly efficient synthetic gRNAs targeting exon 3, which encodes the functional helical cytokine domain in Clcf1 (Morris et al., 2018). The efficiency of *clcf1* target site disruption was confirmed by showing resistance of the targeted DNA sequence to restriction enzyme digest via RFLP analysis and by showing reduced mRNA expression in qRT-PCR (Supplementary Figure 7A, B). Somatic *clcf1* mutants showed no deviations in body length or eye diameter from control gRNA-injected larvae, suggesting that Clcf1 was not essential for general development (Supplementary Figure 7C-F). To create stable germline transmitted mutants, two gRNAs targeting exon 3 were used, leading to a 105-bp deletion in exon 3 corresponding to the gRNA target sites, which resulted in a frameshift that led to a premature stop codon (Supplementary Figures 7G-I). The predicted protein encoded by this allele was truncated at the C-terminal lacking the functional domain (four helical cytokine domain), likely creating a loss-of-function mutant (Supplementary Figure 7I).

To dissect the likely mode of action of Clcf1 signalling, we assessed the impact of *clcf1* disruption on the generation of motor neurons. We assessed cumulative numbers of EdU-incorporating motor neurons (Figure 6A). In unlesioned somatic *clcf1* mutants, ongoing developmental neurogenesis was unaffected by gRNA injection, suggesting a lesion-specific effect of Clcf1 (Figure 6B). The lack of an effect in unlesioned animals was also consistent with the absence of detectable *clcf1* expression without a lesion. Following spinal cord injury, the lesion-induced elevated number of EdU^+^/*mnx1:GFP^+^* neurons was reduced by 31% in *clcf1* somatic mutants compared to larvae injected with control gRNAs (Figure 6B), suggesting that Clcf1 is specifically required for regenerative neurogenesis. Similarly, the lesioned *clcf1* germline mutant displayed a 38% reduction in the number of EdU^+^/*mnx1:GFP^+^* neurons compared to lesioned wild-type siblings (Figure 6C, D) Next, we determined the effect of *clcf1* disruption on acute proliferation of ERGs in lesioned larvae (Figure 6E, F). Unlesioned larvae were not included in the analysis because *clcf1* disruption did not significantly affect motor neuron generation under developmental conditions, suggesting that its role is specific to lesion-induced neurogenesis. We performed a single EdU pulse followed by a 4 h delay using the *her4.1:eGFP* transgenic line in somatic *clcf1* mutants at 24 hpl (Figure 6E). We found that the number of EdU^+^/*her4.1:eGFP^+^* ERGs was reduced by 42 % in *clcf1* somatic mutants compared to control injected larvae, suggesting that Clcf1 promotes progenitor proliferation during regenerative neurogenesis (Figure 6F).

**Fig. 6.**
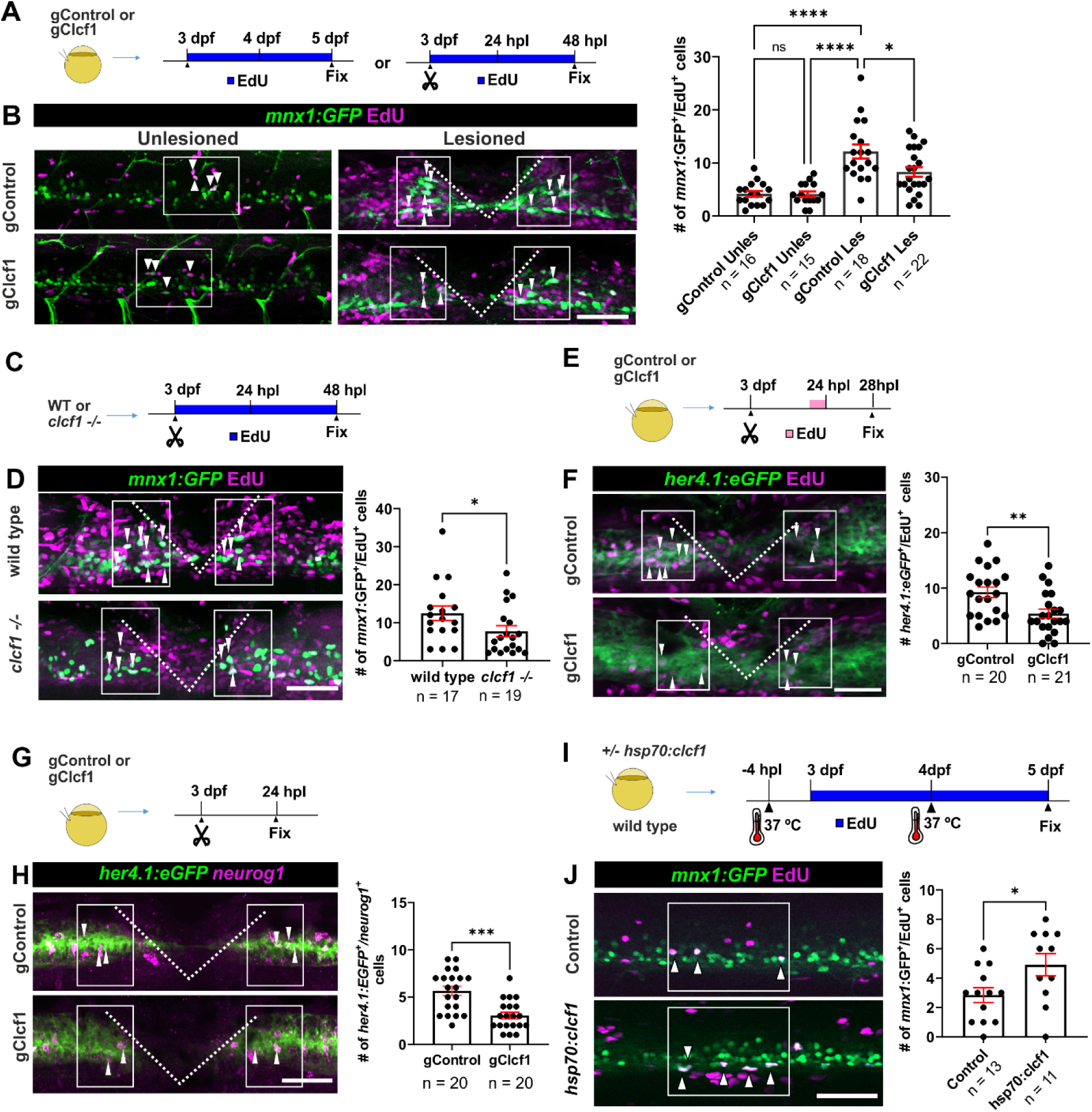
Clcf1 positively regulates motor neuron generation. (**A-B)** (A) shows a schematic of the experiment. (B) Ongoing developmental generation of motor neurons (*mnx1:GFP*+/EdU+) is unchanged in unlesioned somatic mutants for *clcf1*, but lesion-induced generation of motor neurons is decreased in somatic mutants for *clcf1* (gControl Unles: 4.2 cells per larva ± 0.5; gClcf1 Unles: 4.1 cells per larva ± 0.5; gControl Les: 12.1 cells per larva ± 1.3; gClcf1 Les: 8.3 cells per larva ± 0.9; Two-way ANOVA: p < 0.0001; Holm-Šidák’s multiple comparison test: p**** < 0.0001, p* = 0.0415). **(C-D)** (C) shows schematic of the experiment. (D) Lesion-induced generation of motor neurons (*mnx1:GFP*+/EdU+) is decreased in *clcf1* germline stable mutants (wildtype: 12.4 cells per larva ± 1.9; *clcf1 -/-*: 7.7 cells per larva ± 1.5; Mann-Whitney U-test: p = 0.0326). **(E-F)** Lesion-induced proliferation of ERGs (*her4.1:eGFP+/EdU+* pulse) is decreased in somatic mutants for *clcf1* (gControl: 9.2 cells per larva ± 0.9; gClcf1: 5.4 cells per larva ± 0.8; Unpaired t-test: p = 0.0040). (**G-H)** (G) shows a schematic of the experiment. (H) Lesion-induced neurogenic progenitors (*her4.1:eGFP+/neurog1+)* are decreased in number in somatic mutants for *clcf1* at 24 hpl (gControl: 5.6 cells per larva ± 0.5; gClcf1: 3.0 cells per larva ± 0.3; Unpaired t-test: p = 0.0001). **(I-J)** (I) shows schematic of the experiment. (J) Ongoing developmental generation of motor neurons (*mnx1:GFP*+/EdU+) is increased in unlesioned larvae by over-expression of *clcf1* (Control: 2.8 cells per larva ± 0.5; *hsp70l:clcf1*: 4.9 cells per larva ± 0.7; Unpaired t-test: p = 0.0293). **Scale bars:** 50 µm. Dorsal is up and rostral is left.

To assess any changes in the number of progenitor cells undergoing neuronal differentiation, we assessed the number of *neurog1*-expressing neurogenic precursors in *clcf1* somatic mutants following spinal cord injury at 24 hpl (Figure 6G). We found a 46% decrease in the number of *neurog1^+^*/*her4.1:eGFP^+^*ERGs in *clcf1* somatic mutants compared to control larvae (Figure 6H). This reduction was in line with the magnitude of the reduction in the proportion of acutely proliferating ERGs (42%, see above) and thus the reduced number of differentiating ERGs could be a consequence of reduced proliferation in ERGs (Figure 6H).

To determine whether *clcf1* alone could augment neurogenesis, we used injection of an expression plasmid coding for heat-shock inducible *clcf1* expression (*hsp70l:clcf1;* Supplementary Figure 8A,B). A single heat-shock led to robust and stable > 100-fold over-expression of *clcf1* transcript for at least 24 h after heat-shock, as indicated by qRT-PCR (Supplementary Figure 8D, E). Over-expression was also detectable by HCR-FISH in the trunk region in the vicinity of the lesion (Supplementary Figures 8C). For cell counting experiments, we applied two heat-shocks. One 4 hours before the lesion timepoint at 3 dpf and 24 h later to ensure elevated *clcf1* levels throughout the experiment (Figure 6I).

In unlesioned animals, cumulative labelling of motor neurons (*mnx1:GFP^+^*) at 5 dpf (corresponding to 48 hpl in lesioned animals) that incorporated EdU (EdU present from 4 hours before the 3 dpf timepoint to 5 dpf), we observed a 75 % increase in in the number of EdU-labelled motor neurons (Figure 6I and J). In lesioned larvae, no additional generation of motor neurons was observed at 48 hpl after *clcf1* over-expression (Supplementary Figures 8F-H). These observations indicate that *clcf1* over-expression is sufficient to enhance neurogenesis without additional lesion signals in the unlesioned situation. The lack of an additional effect on regenerative neurogenesis after lesion suggests that the lesion-induced expression of *clcf1* already produced maximal effects in neurogenesis that could not be further augmented by exogenous signal. This also suggests that the artificially augmented *clcf1* transcripts (> 100-fold) did not produce ectopic effects driven by high ligand levels alone.

### Il6st is necessary for regenerative neurogenesis

Clcf1 signals through various receptor complexes, all of which include Il6st (Senaldi et al., 1999). To confirm the involvement of Il6st signaling in regenerative neurogenesis, we examined whether Il6st signaling is essential for this process. Our scRNA-Seq analysis revealed widespread expression of *il6st*, including in domain-specific progenitors and highly proliferative ERGs (Figure 7A). Indeed, HCR-FISH analysis of *il6st* expression in spinal cord-lesioned *olig2:GFP* transgenic reporter larvae at 24 hpl showed expression in the ventricular zone, including in the pMN-like *olig2:GFP^+^* cells at 24 hpl (Figure 7B).

**Fig. 7.**
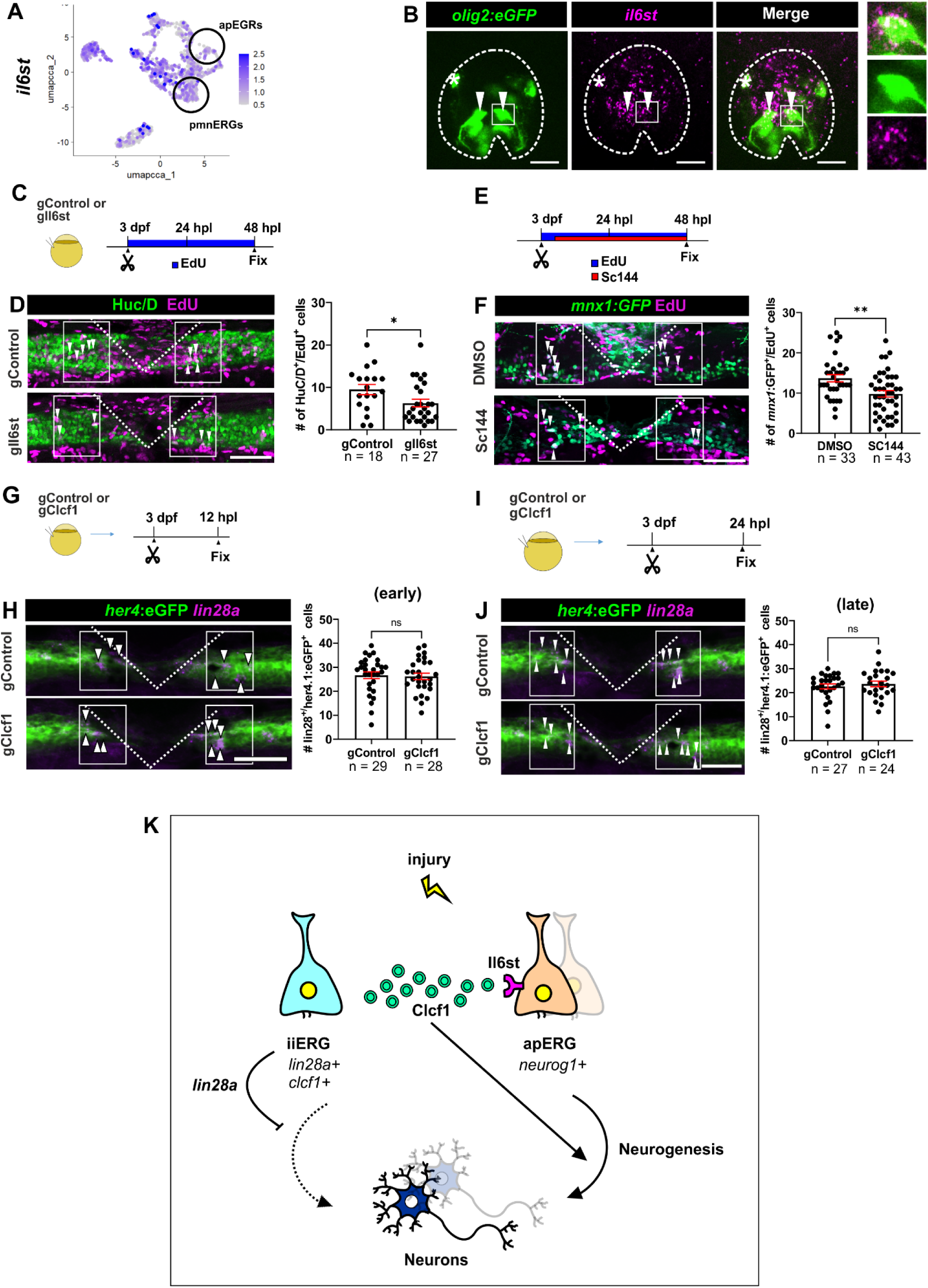
Disruption of Clcf1 receptor Il6st phenocopies the effect observed with *clcf1* disruption. **(A)** Feature plot showing *il6st* expression in all ERGs including motor neuron progenitors (circles). **(B)** HCR-FISH against *il6st* in a cross section of an *olig2:eGFP* animal shows expression in motor neuron progenitors (arrows). **(C-D)** (C) shows schematic of the experiment. (D) Lesion-induced generation of new neurons (HuC/D+/EdU+) is decreased in somatic mutants for *il6st* (gControl: 9.5 cells per larva ± 1.2; gIl6st: 6.3 cells per larva ± 0.9; Mann-Whitney U-test: p = 0.0429). **(E-F)** (E) shows schematic of the experiment. (F) Lesion-induced generation of motor neurons (*mnx1:GFP* +/EdU+) is decreased in Sc144 drug-treated larvae compared to lesioned DMSO-treated animals (DMSO: 13.7 cells per larva ± 0.9; Sc144: 9.8 cells per larva ± 0.8; Unpaired t-test: p = 0.0020). **(G-H)** (G) shows schematic of the experiment. (H) HCR against *lin28a* in somatic mutants for *clcf1* shows no differences in the number of *her4:eGFP*/*lin28*+ cells at 12 hpl (gControl: 26.7 cells per larva ± 1.4; gClcf1: 26.21 cells per larva ± 1.3; Mann-Whitney U-test : p =0.5759). **(I-J)** (I) shows schematic of the experiment. (J) HCR against *lin28a* in somatic mutants for *clcf1* shows no differences in the number of *her4:eGFP*/*lin28*+ cells at 24 hpl (gControl: 22.6 cells per larva ± 1.1; gClcf1: 23.6 cells per larva ± 1.2; Mann-Whitney U-test : p =0.7606). **(K)** Summary schematic showing the proposed role of iiERGs in preserving stemness via *lin28a* and promoting neurogenesis via *clcf1*. **Scale bars:** 50 µm (D, F, H, J), 15 µm (B). Dorsal is up and rostral is left.

To disrupt Il6st signaling, we used highly efficient gRNAs targeting of the *il6st* gene (Supplementary Figure 9), as well as Sc144, a pharmacological inhibitor of Il6st downstream signaling. As a read-out, we used cumulative EdU labelling at 48 hpl as above (Figure 7C, E). We focused only on lesion-induced neurogenesis to specifically address the regenerative role of Clcf1/Il6st signaling and excluded unlesioned larvae from the analysis because disruption of the Il6st ligand Clcf1 selectively impairs regenerative neurogenesis but not ongoing developmental neurogenesis. In lesioned somatic *il6st* mutants, we used HuC/D as a pan-neuronal marker (Figure 7C). This indicated a 34% reduction in the number of newly generated neurons when *il6st* was disrupted (EdU^+^/HuC/D^+^; Figure 7D). Similarly, after Sc144 treatment in transgenic *mnx1:GFP* larvae, we observed a 29 % reduction in the number of newly generated motor neurons (EdU^+^/*mnx1:GFP*^+^; Figure 7E, F) following spinal cord injury, compared to control larvae. These reductions were similar to those observed after *clcf1* disruption, consistent with active Clcf1 signalling during regenerative neurogenesis.

### Clcf1 acts on neurogenesis independently of Lin28a

The observation that *clcf1* overexpression increased neurogenesis in the absence of a lesion made us wonder whether it would also induce the iiERG state. To answer that question, we used HCR-FISH detection of *lin28a*, as a marker for iiERGs, in unlesioned *her4.1:GFP* transgenic animals that were injected with the heat shock-inducible *clcf1* over-expression plasmid. However, we hardly detected any *lin28a^+^/her4.1:GFP^+^*cells in non-heat shocked control, or in the experimental group at 12 h after heat shock (Supplementary Figure 8I, J). This suggested that *clcf1* promotes neurogenesis without a need for *lin28a* expression or an iiERG state.

To determine whether Clcf1 has a role in the maintenance of the iiERG state after lesion, we analysed *lin28a* expression in lesioned somatic *clfc1* mutants, which showed reduced numbers of newly generated motor neurons and *her4.1:GFP^+^* progenitors. We analyzed *lin28a* expression by HCR-FISH at 12 hpl (early), when the number of *lin28a^+^* iiERGs peaked and at 24 hpl (late), when neurogenesis peaked in controls (Figure 7G, I). At both time points, we did not observe any changes in the numbers of *lin28a^+^/her4.1:GFP^+^*iiERGs compared to control guide-injected animals (Figure 7G-J). This suggested that the *lin28a*+ iiERGs state was independent of *clcf1* expression.

Overall, our data support a model in which the iiERG state is induced by spinal cord lesion. Lin28a attenuates neurogenesis from iiERGs while promoting neurogenesis from ERGs via growth factor secretion, e.g. Clcf1 (Figure 7K).

## DISCUSSION

In this study, we provide evidence of the existence of a unique injury-induced population of ERGs during spinal cord regeneration of zebrafish that we named iiERGs. iiERGs are characterized by a stem cell-like transcriptional profile and by the expression of secreted growth factors. These cells emerge prior to the onset of regenerative neurogenesis and are present during the regenerative process, and therefore are in a key position to condition the ERG niche in the lesioned spinal cord.

### A lesion-specific transcriptional profile is induced in a subpopulation of ERGs prior to the onset of regenerative neurogenesis

We identified a lesion-induced transcriptional state in ERGs. This subset of ERGs (iiERGs) is characterized by the upregulation of genes typically associated with stemness, such as *lin28a*, as well as cytokine genes, including *clcf1*, and strongly differs in several key aspects from the neurogenic populations observed during development. Notably, the appearance of iiERGs is temporally confined to the first 12 hours following spinal cord injury, coinciding with an observed delay before the onset of regenerative neurogenesis. We found that regenerative neurogenesis was preceded by a ∼16-hour delay, during which low level neurogenesis was comparable to the ongoing developmental neurogenesis seen in age-matched unlesioned controls. Therefore, we delineated a phase of regenerative neurogenesis starting at approximately 16 hpl. Similarly, during axolotl spinal cord regeneration, a lag phase of 4 days before neurogenesis started was observed (Cura Costa et al., 2021; Rodrigo Albors et al., 2015; Rost et al., 2016). In adult zebrafish telencephalon, a similar lag phase of 3 days was observed before the onset of neurogenesis (Kizil et al., 2012). This reinforces the concept that a lag phase precedes the initiation of regenerative neurogenesis in both species, highlighting a conserved, delayed response of ERGs to spinal cord injury. This pattern suggests that iiERGs represent an early, injury-specific response in the progenitor population, preceding the transition into regenerative neurogenesis and possibly contributing to later stages of regeneration.

The emergence of a specific injury-induced progenitor state has been documented in the adult axolotl telencephalon and zebrafish retina (Fausett & Goldman, 2006; Lust et al., 2022). Following stab injury in the axolotl telencephalon, ERGs upregulate regeneration-specific genes, such as *Plat, Anxa2, Kazald1, Erbb3, Col7a1* and *Runx1*, prior to the onset of neurogenesis (Lust et al., 2022). Similarly, retinal injury activates an injury-specific program in Müller glia (MG), which are typically non-neurogenic and non-proliferative under homeostatic conditions, reprogramming them to re-enter a neurogenic state and generate new photoreceptors (Bernardos et al., 2007; Fausett & Goldman, 2006; Fimbel et al., 2007; Raymond et al., 2006). Hence, de-differentiation of progenitors may be a hallmark of regenerating systems.

### iiERGs lack domain-specific transcriptional programs and adopt a neural stem cell like state

In both larval and adult zebrafish spinal cords, ERGs retain their dorso-ventral progenitor identity from development as evidenced by the expression of dorso-ventral specific markers in ERGs in lesioned and unlesioned spinal cords (Briona et al., 2015; Kuscha et al., 2012a; Reimer et al., 2008). Consistent with this, our 12 hpl larval scRNA-Seq dataset identifies ERG populations corresponding to distinct dorso-ventral progenitor domains in both unlesioned and lesioned animals. Moreover, following spinal cord lesion, we observe an increase in the number of EdU-labeled proliferative cells as well as in *neurog1^+^* neurogenic *olig2^+^* pMN domain progenitors by HCR-FISH, indicating that domain-specific progenitors acquire neurogenic competence in response to a lesion. Similarly, in adult zebrafish, ventral *olig2^+^* pMN progenitors re-enter the cell cycle and give rise to regenerated motor neurons after spinal cord lesion (Reimer et al., 2008). Together, these findings establish that regenerative neurogenesis likely originates from domain-specific progenitor populations. Interestingly, scRNA-seq analysis shows that iiERGs hardly express domain-specific marker genes. The lack of domain-specific progenitor properties may reflect a de-differentiation of iiERGs toward a less committed neural stem cell-like state. iiERGs express several genes that are associated with maintaining stem cell identity, such as *lin28a*, *sox2* and *mycb*, suggesting that iiERGs may adopt a stem cell like state. (Moss et al., 1997; Moss & Tang, 2003; Ouchi et al., 2014; Yang & Moss, 2003; Yu et al., 2007). For example, *lin28a* is expressed in pluripotent cells and stem cell-like cells, and not in differentiated tissues (Moss et al., 1997; Moss & Tang, 2003; Ouchi et al., 2014; Yang & Moss, 2003; Yu et al., 2007). Lin28a regulates pluripotency and self-renewal in embryonic stem cells and is capable of reprogramming somatic cells to a pluripotent state (Bin et al., 2012; Hanna et al., 2009; Yu et al., 2007). Additionally, iiERGs express the stemness genes *sox2*, *klf4* and *mycb*, that are also associated with pluripotency and stem cell renewal (An et al., 2019). More specifically, iiERGs showed enriched expression of neural stem cell-related genes, including *numb*, *numbl*, and *her9* (Engel-Pizcueta et al., 2025; Knoblich et al., 1995). This aligns with the notion that iiERGs adopt specific characteristics of neural stem cells. Therefore, the expression of *lin28a* and other stemness genes aligns with iiERGs adopting characteristics of stem cells. Similarly, Müller glia in the injured zebrafish retina also upregulate the stem cell-related genes *lin28a*, *sox2*, *klf4,* and *cmycb,* and additionally also upregulate *pou5f1*/*oct4* during de-differentiation, which was not detectable in iiERGs (Gorsuch et al., 2017; Mitra et al., 2019; Ramachandran et al., 2010; Sharma et al., 2019). Hence, iiERGs in the spinal cord likely differ from Müller glia in their potential.

Another argument for stemness of iiERGs is their lack of expression of pro-neural genes that are typically involved in neural differentiation and are expressed in other ERG populations. For example, *neurod4, atoh1b, neurog1* and *neurod1* are not expressed in iiERGs, but in proliferating ERGs (Bertrand et al., 2002; Castro et al., 2011). The absence of pro-neural genes is a key characteristic of neural stem cells in various systems (Bertrand et al., 2002; Ross et al., 2003). Consistently, upregulation of neural stem cell markers, alongside downregulation of pro-neurogenic genes, have been reported in spinal ERGs of frogs and axolotl after lesion (Rodrigo Albors et al., 2015; Walder et al., 2003). This suggests that iiERGs, like neural stem cells, may maintain an undifferentiated state during the early phases of regeneration, preserving their potential to self-renew and perhaps differentiate at later time points.

### Lin28a promotes maintenance of iiERGs and apERGs

Consistent with a role in maintaining progenitors during regenerative neurogenesis, we found that *lin28a* depletion after spinal cord lesion reduces the number of *clcf1+* iiERGs and proliferating ERGs, while increasing the number of differentiated neurons. This is comparable to the genetic deletion of other stem cell-related factors in adult neural stem cells. For instance, deleting the stemness-associated transcription factor ZEB1 in activated radial glia of the adult mouse hippocampus leads to an increase in the number of newborn neurons (Gupta et al., 2021). Moreover, our findings show that *lin28a* is essential for maintaining the ERGs after spinal cord lesion. Deletion of *lin28a* results in a decrease of both iiERGs, which coincides with increased pro-neural commitment and differentiation of ERGs. Similar to our observation, Lin28a is highly expressed in cortical neural progenitors during mouse development and is required to maintain a neural progenitor pool and to prevent premature neuronal differentiation (Yang et al., 2015). Interestingly, the combined deletion of Lin28a and the paralogue Lin28b in the mammalian neuroepithelium results in a decreased number of Olig2^+^ progenitors and increased number of progenitors committed to a motor neuron fate in the posterior neuroepithelium (Herrlinger et al., 2019). In the zebrafish retina, the function of *lin28a* is mainly discussed as a re-programming factor for these facultative progenitor cells. In iiERGs, which are closely related to developmental ERG progenitors and may therefore not need extensive re-programming, our findings uncover a previous unrecognized link between lesion-specific *lin28a* expression and the maintenance of progenitors during regenerative neurogenesis and suggest that Lin28a might balance ERG self-renewal and neuronal differentiation during spinal cord regeneration. The observation that after disruption of *lin28a* also the number of actively proliferating ERGs is reduced, is likely linked to the reduced expression of *clcf1* after *lin28a* disruption, since *clcf1* acts as a growth factor that stimulates ERG proliferation.

### Clcf1/Il6st signaling may contribute to the neurogenic niche

Growth factors and cytokines play a crucial role in promoting regenerative neurogenesis (Cavone et al., 2021). Notably, iiERGs upregulate several cytokine and growth factor-related genes, which may condition the spinal progenitor niche to induce regenerative neurogenesis. Disruption of the cytokine Clcf1 and its receptor Il6st impairs regenerative neurogenesis, reducing both proliferation and neural commitment. These findings suggest that Clcf1 signaling supports regenerative neurogenesis. In this model, Clcf1 secreted by neighboring iiERGs may act on the neurogenic and proliferative ERG population via activation of the Il6-signaling pathway in the spinal cord, as indicated by the expression and functional importance of the Il6st receptor in these cells.

Interestingly, cytokines of the IL-6 family, including Clcf1, have been shown to enhance MG reprograming and proliferation after retinal injury in adult zebrafish (Boyd et al., 2023; Zhao et al., 2014), although whether this leads to enhanced neurogenesis has not been analyzed.

### Clcf1 does not induce the iiERG state

We show that disruption of *clcf1* does not affect the emergence of *lin28a*-expressing iiERGs in the lesioned spinal cord. Moreover, overexpression of *clcf1* in the unlesioned spinal cord stimulates ongoing developmental neurogenesis, independent of *lin28a* upregulation in ERGs. These findings support a model in which Clcf1 can promote regenerative neurogenesis independently of the induction of an iiERG state in the spinal cord. In contrast, in retinal Müller glia, Il6-signaling mediated by Il11 and Clcf1 is necessary for the induction of *lin28a* and other reprograming genes, such as *ascl1a* after retinal injury indicating that Clcf1 is already required for adopting the activated MG state (Zhao et al., 2014).

Our finding, that *clcf1* overexpression enhances ongoing neurogenesis in the unlesioned spinal cord, suggests that spinal ERGs are already responsive to Clcf1-mediated signaling even before injury, and can initiate enhanced proliferation in response to this signal. In contrast, *clcf1* overexpression in lesioned larvae did not further increase neuron generation during regenerative neurogenesis, indicating that endogenous levels were already maximally effective. In mammals including humans, the Clcf1 related Clrf1/Il6st receptor ligand CNTF has been shown to improve neuronal survival in the CNS (Rhee et al., 2013; Beurrier et al., 2010; Jeong et al., 2015). Interestingly, in mammals, including humans, ERGs appear to be sensitive to CNTF (Leibinger et al., 2009; Rhee et al., 2013; Wang et al., 2020; Beurrier et al., 2010; Jeong et al., 2015), indicating that mammalian spinal progenitor cells might also be responsive to Clcf1. This suggests potential strategies for reintroducing neurogenic potential into spinal progenitors of non-regenerative species.

In summary, our study provides evidence that a unique progenitor cell state of iiERGs is induced by spinal cord injury in zebrafish. These iiERGs adopt neural stem cell-like characteristics and upregulate key stemness factors such as *lin28a*, along with expression of growth factors like *clcf1*. Our data suggest that iiERGs maintain a stem cell-like state to maintain the progenitor pool in two ways, by preventing neuronal differentiation and promoting replenishment of progenitors by stimulating their proliferation.

## MATERIALS AND METHODS

### Animals

All zebrafish lines were raised and kept under standard conditions (Westerfield, 2000). Zebrafish experiments were performed under state of Saxony licenses TVV 36/2021, TVV 45/2018, and holding licenses DD24-5131/364/11, DD24-5131/364/12. For experimental conditions, larvae up to an age of 5 days post-fertilization (dpf) were used of the following lines: wild type (AB); Tg(*mnx1:GFP*)^ml2tg^, abbreviated as *mnx1:GFP* (Flanagan-Steet et al., 2005); Tg(*her4.1:eGFP*)^y83Tg^, abbreviated as *her4.1:eGFP*) (Yeo et al., 2007); Tg(*olig2:EGFP*), abbreviated as *olig2:EGFP* (Shin et al., 2003); and Tg(*pdgfrb:Gal4ff*)^ncv24^; Tg(*UAS:GFP*), abbreviated as *pdgfrb:GFP* (Ando et al., 2016).

### Spinal cord lesion

Lesions of larvae were performed as previously described (Ohnmacht et al., 2016). Briefly, at 3 dpf, zebrafish larvae were deeply anaesthetized in E3 medium (Nüsslein-Volhard & Dahm, 2002) containing 0,02 % of MS-222 (Sigma). Larvae were then transferred to an agarose plate. After removal of excess water, larvae were placed in lateral positions. A transection of the entire spinal cord was made using a 30G syringe needle at the level of the 15^th^ myotome, without injuring the notochord.

### Generation of somatic mutants

Somatic mutations were generated mainly following a previous protocol (Keatinge et al., 2021). CRISPR gRNAs were selected using the CRISPRscan website (www.crisprscan.org/gene) webtools. gRNAs were chosen based on the CRISPRScan score (preferentially > 60), and on the availability of a suitable restriction enzyme recognition site (www.snapgene.com) that would be destroyed by the gRNA. Additional criteria were to avoid the first exon and to target functional domains. 1 nL of gRNA mix (1 µl of Tracer (5nM, TRACRRNA05N, Sigma-Aldrich), 1 µl of gRNA (20µM, Sigma-Aldrich), 1 µL of phenol red (P0290, Sigma-Aldrich) and 1 µL of Cas9 (20µM, M0386M, New England Biolabs)) was injected into one-cell stage embryos. gRNA efficiency was tested by restriction fragment length polymorphism analysis (RFLP) in a proportion of larvae for each experiment as an internal experimental control (Keatinge et al., 2021). Sequences for gRNAs and primers for RFLPs can be found in Suppl. Table 1. All experiments conducted with somatic mutants were carried out with a control injected group with a non-targeting gRNA.

### Generation of *clcf1* germline mutant

The *clcf1* mutant stable line was generated by raising somatic mutants as potential founders, as described (Keatinge et al., 2021). The founder selected to generate the stable line carried a 105 bp deletion in exon 3 that generated a frame shift with a premature stop codon. Sequencing primers were the same as for RFLP.

### Single-cell sequencing

We performed cell dissociation as described previously (Docampo-Seara et al., 2026). In brief, we cut trunk around the injury site from 200 larvae for each condition using Tg(*her4:EGFP*) line. We sorted 125,000 GFP cells using FAC-sorting. The cells were either directly used for single-cell sequencing or used for nuclei extraction.

For single-cell sequencing, we directly sorted cells in 1x PBS with 0.4 BSA with final concentration of BSA to be between 0.2-0.8%. The cells were pelleted at 500g for 5 mins at +4 °C, supernatant was removed to keep around 20µL volume. Cells were pipetted and 1µL of the cells were diluted in 9µL of trypan blue. Viable cells were counted under a light microscope using a Neubauer chamber. Samples with over 90% viability were used for encapsulation. The encapsulation and single-cell sequencing were done as described previously (Docampo-Seara et al., 2026). Genome alignment was also performed as described previously (Docampo-Seara et al., 2026).

### Single-cell analyses

For single-cell analyses, we used Seurat V5 as described previously (Docampo-Seara et al., 2026). In brief, we used the filtered matrices from Cellranger outputs. The following cells were kept for analyses; (i) cells with more than 200 genes, (ii) with fewer than 25000 total reads, (iii) less than 20% mitochondrial genes, (iv) less than 10 fold difference between total reads per cells and number of genes detected per cells, (v) more than 1.3 fold ratio between total reads and number of genes detected per cells.

Additionally, genes found in fewer than 5 cells were removed from analyses. The integration was done using the first 30 PCAs, number of dimension 30. Following normalization, all genes genes were used for scaling by regressing out nCount_RNA (total reads). Dimensional reduction was done using UMAP and cell clusters were identified by using resolution 1.

### Cell type annotation

We used the following genes to identify first main cell types; (i) ERG cells; *gfap, fabp7a*, (ii) inhibitory neurons (*gad2, gad1b*), (iii) Glycinergic Neurons (*slc6a5*), (iv) Excitatory Neurons (*slc17a6a, slc17a6b*), (v) motor neurons (*chata, mnx1*), (vi) Intraspinal serotonergic neurons (ISN) (*thp2, fev*), (vii) CSF-cNs (*sst1.1* and *urp1*), (viii) Oligodendrocyte Precursor Cells and ODs (*aplnra, cldnk*), (ix) Fibroblast Cells (*pdgfrb, col12a1a, col12a1b*), (x) Enteric Nervous System (*phox2bb, phox2a*), (xi) Schwann Cells (*slc35c2*), neural progenitor ERGs (nERGs) (*nhlh2, nfic, nfia*), Lateral Floor Plate (*nkx2.9*), Floor Plate (*shha*), Roof Plate (*mdka*), Rohon Beard (*hpca*). The clusters that were not annotated were named with their cluster numbers (e.g., c#34).

### Sub-clustering of ERG cells

For sub-clustering ERG cells, we used LFP, ERG cells and nERG clusters for integration using the above Seurat pipeline, with exception of using 10 PCA and 10 dims. The annotation of the sub-cluster of ERGs was done using top marker genes and searching for domain markers. In brief, we used the following markers; (i) *zic1, zic2a, olig3, msx1a* and a broad dorsal marker *pax7a* for a mix of populations that we named as dorsal ERGs (dERGs), (ii) *olig2* for pMN domain, (iii) *pcna, mki67* for actively proliferating ERGs (apERGs), (iv) *nhlh2, nfic, nfia* for neural progenitor ERGs (nERGs), (v) *sv2a, chata, mnx1* for commited MN ERGs, (vi) clcf1 for iiERG, (vii) *dbx1*a for p1 domain ERGs (p1ERGs).

### Trajectory inference using scVelo

RNA velocity was estimated using scVelo (v0.3.4) with the dynamical model (Bergen et al., 2020). Splicing kinetics were recovered for all genes passing standard filtering using scv.tl.recover_dynamics, and per-gene model fit quality was quantified by the likelihood score (fit_likelihood). To construct the velocity graph using genes with well-resolved splicing kinetics while minimizing noise from poorly fitted genes, we restricted velocity estimation to genes with the highest likelihood values (scv.tl.velocity_graph(adata, gene_subset=top_genes)), following the high-likelihood gene selection strategy described by Furtwängler *et al. (Furtwängler et al., 2025)*. The inferred directions of cell trajectories were robust to the size of the high-likelihood gene set and remained consistent across analyses using the top 100, 300, 500, and 800 genes; the top 500 genes were used for visualization of velocity stream plots projected onto the UMAP embedding.

### *In situ* hybridization chain reaction (HCR-FISH)

Larvae were collected in E3 medium and incubated with 0.003 % 1-phenyl 2-thiourea (PTU) at 6 hpf to suppress melanocyte development. Larvae were fixed overnight in 4 % paraformaldehyde (PFA) at 4 °C and then transferred to 100 % Methanol (MeOH) at -20 °C overnight for permeabilization. The next day, larvae were rehydrated in a graded series of MeOH at room temperature (RT) and subsequent washes in phosphate buffered saline with 0.1 % Tween (PBST). Thereafter, larvae were treated with 500 μL proteinase K (Roche) at a concentration of 30 μg/mL for 45 min at RT, followed by post-fixation in 4 % PFA for 20 min RT and washes with PBST (3 x 5 min). Pre-hybridisation was performed using 500 μL of pre-warmed probe hybridisation buffer (Molecular Instruments (Choi et al., 2018)) for 30 min at 37 °C. The probe sets, which were designed based on the NCBI sequence by Molecular Instruments (Choi et al., 2018), were prepared using 2 pmol of each probe set in 500 μL of pre-warmed hybridisation buffer. Then, buffer was replaced with probe sets and incubated for 16 h at 37 °C. On the following day, samples were washed 4 x 15 min with pre-warmed probe washing buffer (Molecular Instruments) at 37 °C followed by 2 x 5 min in 5x sodium chloride sodium citrate with 0.1 % Tween (SSCT) at RT. Larvae were then treated with 500 μL of room temperature equilibrated amplification buffer (Molecular Instruments, LA, USA) for 30 min at RT. Hairpin RNA preparation was performed following the manufacturer’s instructions. Samples were incubated with Hairpin RNAs for 16 h in the dark at RT. Finally, samples were washed for 3 x 30 min in 5 x SSCT at RT and immersed in 70 % glycerol for imaging on a Zeiss LSM980 confocal microscope.

### RNAscope

RNAscope fluorescent in situ hybridization was performed using the RNAscope® Multiplex Fluorescent Reagent Kit v2 (Advanced Cell Diagnostics, Bio-Techne) and adapted for whole-mount zebrafish larvae following the manufacturer’s protocol with modifications based on Gross-Thebing et al. (2020). Larvae at 5 days post-fertilization (dpf) were fixed in 4 % paraformaldehyde (PFA) in phosphate-buffered saline (PBS) for 40 minutes at room temperature (RT). Following fixation, larvae were washed three times for 5 minutes each in 0.1 % Tween-20 in PBS (PBST), then dehydrated at RT through a graded methanol (MeOH) series: 25 %, 50 %, 75 %, and 100 % MeOH (5 minutes each). Larvae were stored in 100 % MeOH at −20 °C overnight or longer until further processing. For probe hybridization, larvae were rehydrated through a reverse MeOH series (75 %, 50 %, 25 %, and PBST) and air-dried for 10 minutes at RT, ensuring a thin residual film of MeOH remained to preserve tissue morphology. Larvae were then treated with two drops of Protease Plus solution (ACD Bio) and incubated for 1 hour at 40 °C in a water bath. Probes were warmed to 40 °C for 10 minutes prior to use. Hybridization was carried out by incubating larvae in 50 μL of probe solution mixed with probes in a C1:C2:C3-50:1:1 ratio overnight at 40 °C. The following day, samples were washed in 0.2× SSC with 0.01 % Tween-20 (SSC-Tween) at RT in three sequential steps: 5 minutes, 10 minutes, and 15 minutes. To improve tissue integrity, larvae were post-fixed in 4 % PFA for 10 minutes at RT, followed by another round of SSC-Tween washes.

Signal amplification was performed via sequential incubation in amplification reagents Amp1 (30 minutes at 40 °C), Amp2 (15 minutes at 40 °C), and Amp3 (30 minutes at 40 °C), with SSC-Tween washes in between each step as previously described. For probe detection, the appropriate HRP step was selected based on the probe channel for 15 minutes at 40 °C, washed, and then incubated with TSA Vivid fluorophore (PerkinElmer) diluted 1:1500 in TSA buffer for 30 minutes at 40 °C. Fluorophores used were selected based on the imaging setup and experimental needs. Following signal development, HRP activity was blocked by incubation with HRP blocker for 15 minutes at 40 °C. SSC-Tween buffer and amplification reagents were prepared as per manufacturer instructions. All RNAscope probes, amplification reagents, and detection systems were obtained from Advanced Cell Diagnostics (Bio-Techne, Newark, CA, USA).

### Combining EdU with motor neuron and ERG labelling and birth dating analysis

Labelling newly generated motor neurons followed previous protocols (Ohnmacht et al., 2016). Briefly, lesioned and unlesioned larvae were incubated in E3 medium containing 1 % dimethylsulfoxide (DMSO, Sigma-Aldrich, Taufkirchen, Germany) and 1:100 EdU (Thermo-Fisher, Schwerte, Germany) for 48 h. Samples were fixed in 4 % PFA for 3h at RT and permeabilised in 100 % MeoH at -20 °C for at least overnight. Larvae were then rehydrated, tails were dissected and incubated in proteinase K (Roche, Sigma-Aldrich) for 45 min at RT at a concentration of 10 μg/mL and then postfixed at RT in 4 % PFA for 15 min. Then, larvae were incubated for 20 min at RT in 0.5 % DMSO-PBST and incubated for 2 h at RT with the Click-it Chemistry Edu Alexa 647 kit (Thermo-Fisher, Waltham, USA) as described by the manufacturer. Subsequently, tails were washed in PBST and incubated with blocking buffer solution (2 %BSA, 1 % DMSO, 1 % Triton, 10 % Normal donkey serum, in PBST) for 1 h at RT and then with a chicken anti-GFP antibody, (1:200, Abcam, Cambridge, UK) or the anti-HuC/D antbody (1:50, Invitrogen, California, USA) at 4 °C for 72 hours. Secondary antibody (alexa-488 anti-chicken, 1:200, Jackson Immuno) incubation took place overnight at 4 °C. Finally, samples were washed in PBST and transferred to 70 % glycerol in PBST, mounted and imaged. Imaging was performed using a Widefield Axio Observer (Inverted) with ApoTome 2 Unit (Zeiss). Images were analysed using Fiji (Rueden et al., 2017; Schindelin et al., 2012). Newly generated motor neurons were counted by assessing the number of double-positive cells for GFP and EdU. Cells were counted manually along the Z-stack, ensuring counting individual cells in single optical sections, in 50 µm windows located at both sides of the injury site (100 µm in total). To mitigate bias, experimental conditions were occluded for the analysis. At least two independent experiments were performed. Individual experiments were merged and analysed for statistical significance.

To label proliferating ERGs, we followed the same experimental procedure, imaging and cell counting analysis as for newly generated motor neurons using the ERG reporter line *her4.1:EGFP* instead of the *mnx1:GFP* transgenic line.

For birth dating analysis, the protocol was performed as above, but the EdU incorporation was performed for only for 30 min in 10 µM EdU in 15 % DMSO containing E3. During this process, larvae were kept on ice to slow down cell cycle and to ensure labelling of mitotic cells at the desired time point.

### Tissue sectioning

Larvae stored in 100 % methanol at -20 °C were rehydrated in steps of 75, 50, 25 and 0 % methanol in PBST and fixed using 4% PFA for 20 minutes at room temperature. Following PBST washes, the larvae were mounted in 3-4 % Biozym agarose (Biozym Scientific GmbH, Germany), where care was taken to ensure the agarose was not too hot for the tissue, and the larvae were positioned using insect pins. Then a Leica VT 1200S vibratome (Leica Biosystems, Nussloch, Germany) was used to create spinal cord sections of 4-5 dpf zebrafish larvae, using the settings: 60 μM thickness, 0.20 mm/s speed and amplitude of 0.2. Sections were placed directly into PBS. Larvae were always placed with the head at the top of the agarose so that the proximity to the lesion site could be estimated by the size of the yolk. Sections within approximately 250 μM of the lesion site were selected for HCR-FISH.

### Quantitative RT-PCR

Following spinal cord lesion of 3 dpf, RNA extraction was performed at 1 dpl using the RNeasy Mini Kit (Qiagen) according to the manufacturer’s instructions (40 larvae/condition). The injury site was enriched by cutting away rostral and caudal parts from the larvae at positions approximately 3 somites away from the lesion site in either direction. cDNA was synthesized using the iScript™ cDNA Synthesis Kit (BioRad, cat #. 1708890) according to the manufacturer’s instructions. qRT-PCR was performed using the SsoAdvanced Universal SYBR Green Supermix kit (BioRad) according to the manufacturer’s instructions and run in triplicates on a LightCycler 480SW instrument (Roche). For gene-specific primers see Suppl. Table 2. Each condition was normalized to a housekeeping gene and the experimental group was compared to controls by normalizing to the control group. At least three individual experiments were run. For statistical analysis, a One-Sample T-test was used.

### Drug treatment of larvae

Sc144 (Fisher Scientific) was dissolved in DMSO to a stock concentration of 10 mM. The working concentration was 1 µM prepared by a dilution of the stock solution in E3. Larvae were pre-treated for 2 h before injury and then incubated for 48 hpl. Control larvae were treated with DMSO at the same concentration.

### Generation of heat shock plasmid and treatment

To create the pTol-hsp70l:clcf1-T2a-mcherry plasmid, we modified the pTol-hsp70l:mCherry-T2A-atho1b vector (Ezhkova et al., 2023). We PCR-amplified the *clcf1* coding sequence from genomic cDNA and PCR-amplified T2a-mcherry with specific primers. Then, we performed a simultaneous multiple fragment ligation in the opened backbone vector. The final construct pTol-hsp70l:clcf1-T2a-mcherry was verified by sequencing, qRT-PCR and HCR for over-expression and mCherry fluorescence after injection.

### Bias mitigation and statistical analysis

The experimental conditions were occluded before manual counting and analysis and processing was automated where possible (e. g. measurements of axonal regrowth and intensity measurements). For manual counts of EdU+ cells, pilot experiments were performed by two independent experimenters, both showing comparable effect sizes. Experiments were performed in duplicate, if not indicated differently.

Statistical methods to analyse scRNAseq data are described above. For all other experiments, quantitative data was tested for normality using Shapiro-Wilk’s W-test. Then, parametric (unpaired Student’s t-test or one-way ANOVA) and non-parametric (Mann-Whitney U test or Kruskal-Wallis test) were applied as appropriate. Error bars in figures indicate standard error of the mean (SEM), p-values were represented as asterisks in the figures (*p < 0.05; **p < 0.01; *** p < 0.001) and exact p-values are given in figure legends. To generate graphs and for statistical analysis we used GraphPad Prism 8 (GraphPad Software, Boston, USA).

## Figure preparation

Images were adjusted for brightness, contrast and intensity. Drawings were made using CorelDraw X8 (Corel/Alludo, Ottawa, Canada). Figure plates were prepared using CorelDraw X8 (Corel/Alludo, Ottawa, Canada).

## ACKNOWLEDGEMENTS

We thank Drs Thomas Theil and Anna Poetsch for critically reading the manuscript; Dr. Stefan Hans for reagents; Drs Hella Hartmann and Ruth Hans for imaging advice; and Dr. Judith Konantz, Marika Fischer, and Silvio Kunadt for fish care. This work was supported by the Light Microscopy Facility, the DRESDEN-concept Genome Center, the Flow Cytometry Facility, and the Zebrafish Facility, all core facilities of the Center for Molecular and Cellular Bioengeneering (CMCB) at the Technische Universität (TU) Dresden. Funding was provided by an Alexander von Humboldt Stiftung Professorship award (to CGB), and TU Dresden core funding (to CGB).

## AUTHOR CONTRIBUTIONS (CRediT nomenclature)

**MW:** conceptualization, methodology, validation, formal analysis, investigation, writing - original draft, writing - review and editing, supervision; **RB:** conceptualization, methodology, validation, formal analysis, investigation, supervision; **ADS:** conceptualization, methodology, validation, formal analysis, investigation, writing - original draft, writing - review and editing, visualisation; **MIC**: conceptualization, methodology, formal analysis, investigation; writing - original draft; **KL:** investigation; **MB:** investigation; **SC:** formal analysis, investigation, writing - original draft; **TMT**: investigation; **AB:** investigation; **DZ:** investigation; **RS**: investigation; **LS:** investigation, **DB:** investigation, **TB:** conceptualization, writing - original draft, writing - review and editing, supervision, funding acquisition; **CGB:** conceptualization, writing - original draft, writing - review and editing, supervision, funding acquisition.

## SUPPLEMENTARY FIGURES

**Supp. Fig. 1:**
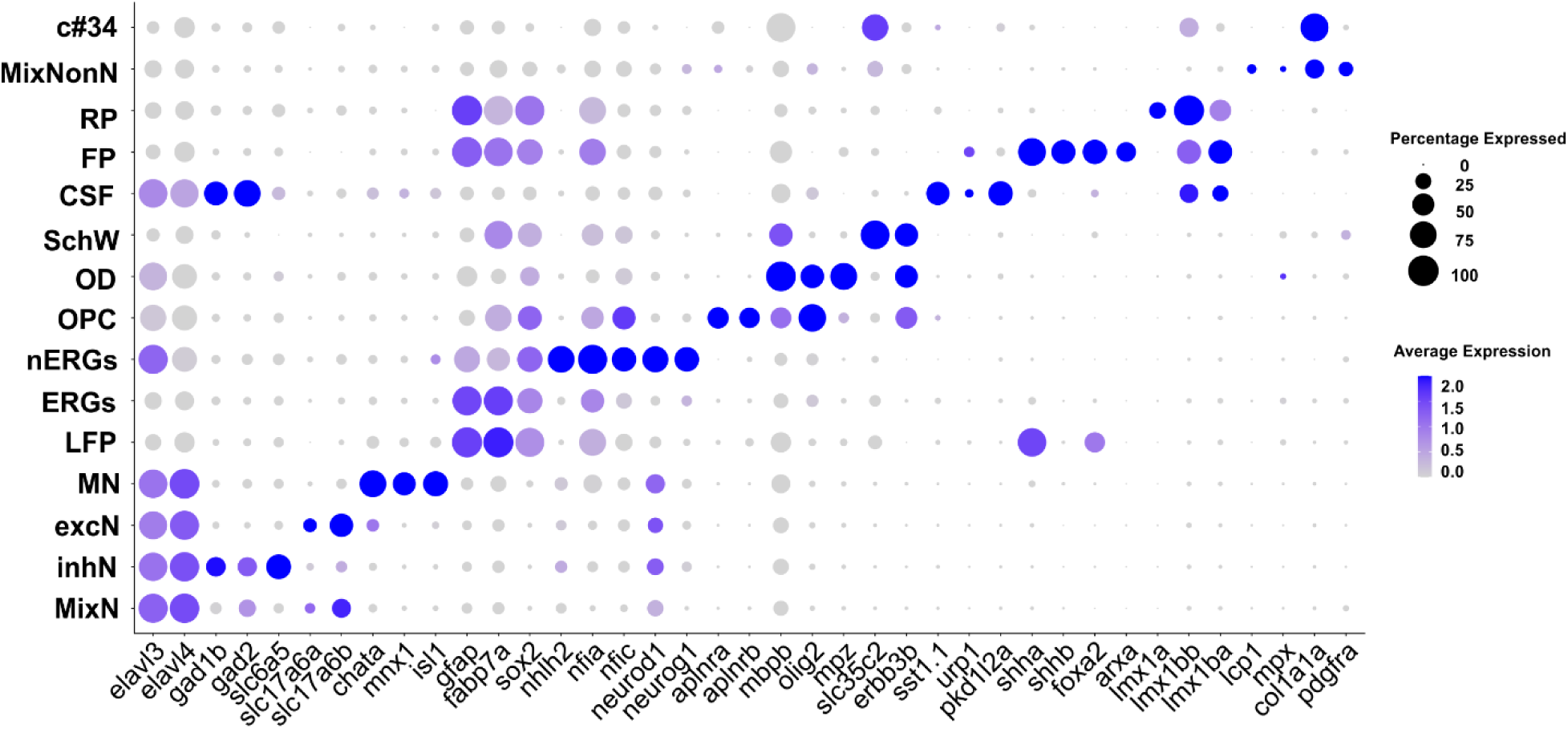
Identity of cell clusters obtained by unsupervised clustering of all cells in sample. Dot plot showing the marker gene expression for the main cell types. MixNonN: Mix of non-neural cells; RP: roof plate; FP: floor plate; SchW: Schwann Cells; OD: oligodendrocytes; OPC: oligodendrocyte precursor cell; nERGs: neural progenitor ERGs; ERG: ependymo-radial glia; LFP: lateral floor plate; MN: motor neuron; excN: excitatory neuron; inhN: inhibitory neuron; MixN: mix of excitatory and inhibitory neurons.

**Supp. Fig. 2.**
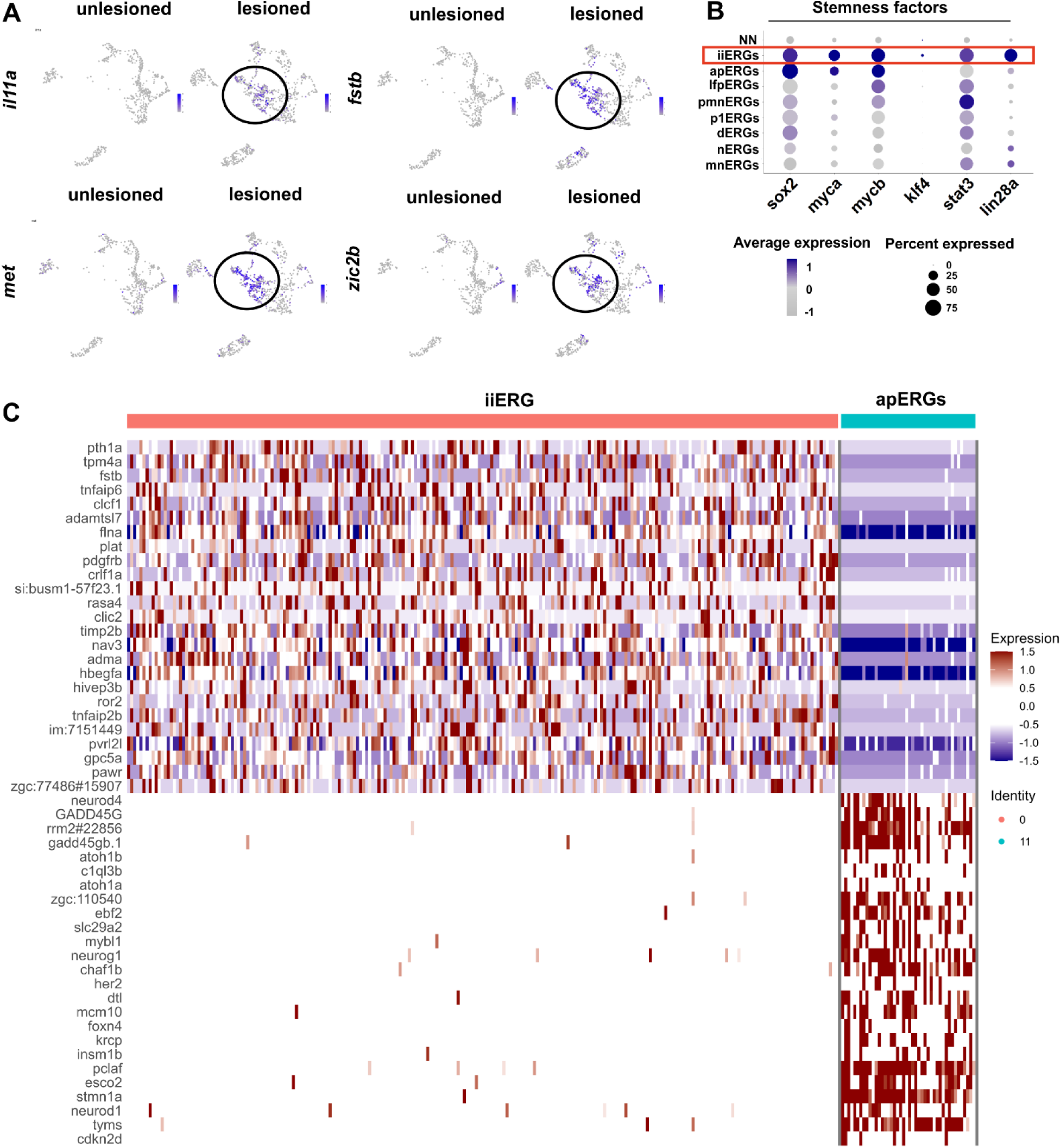
iiERGs express several marker genes. **(A)** Feature plots showing injury-induced specific genes in iiERGs **(B)** Dot plot showing enriched expression of stemness factors in iiERGs. **(C)** Heatmap showing differential expressed genes (DEG) between iiERGs and apERGs. iiERGs express pro-neurogenic signals and Tnf responsive genes, in contrast to apERGs, which express neurogenic genes.

**Suppl. Fig. 3.**
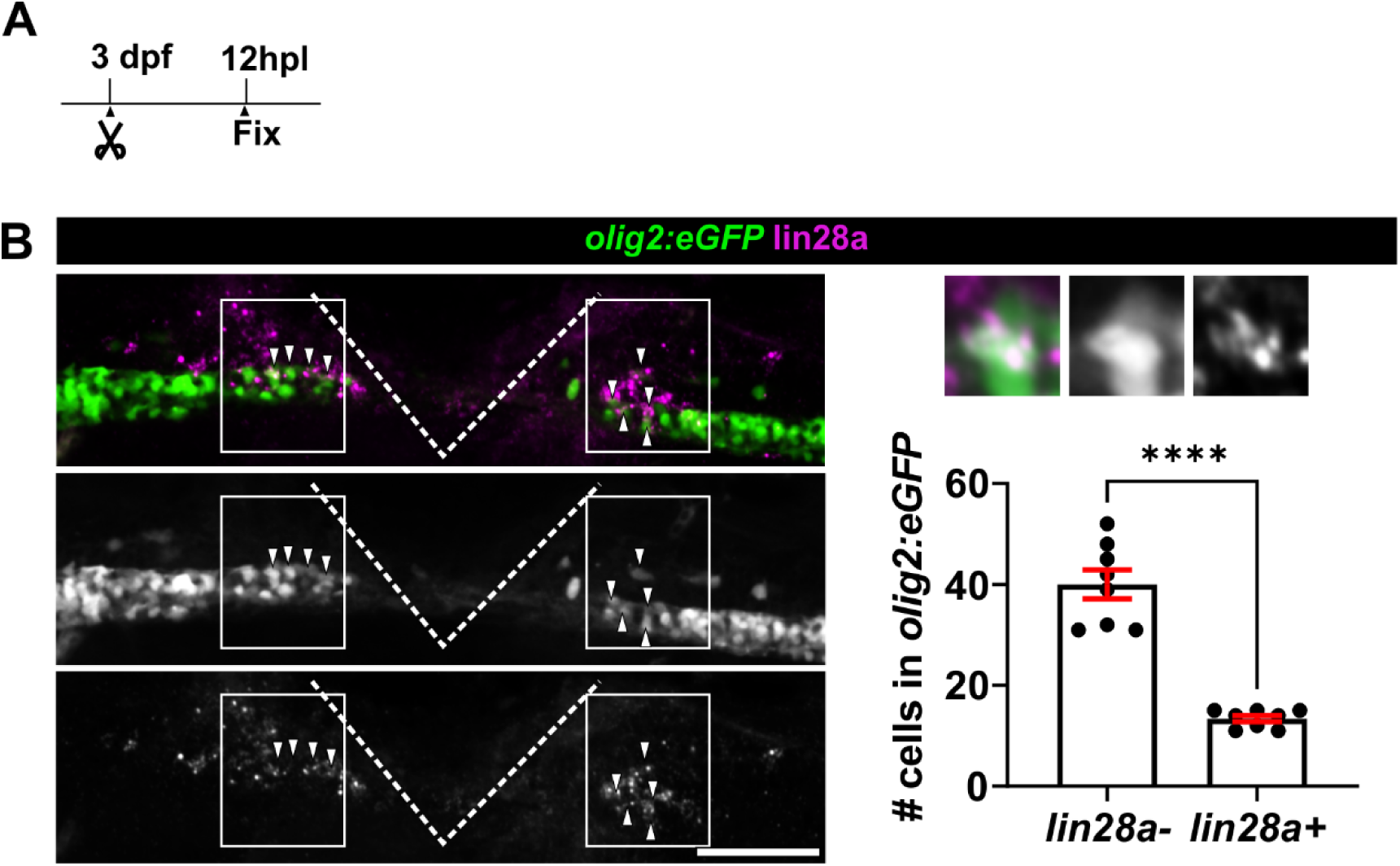
*lin28a* is expressed in *olig2:eGFP*-expressing progenitors after spinal cord lesion. **(A)** HCR-FISH against *lin28a* in *olig2:eGFP* animals at 12 hpl showing double-positive cells. **(B)** Quantification showing that 33.4% of pMN progenitors at the lesion site express *lin28a* (*lin28a-:* 40 ± 2.8 cells; *lin28a+:* 13.4 ± 0.6 cells; Unpaired t-test: p < 0.0001)**. Scale bar:** 100 µm. Dorsal is up and rostral is left.

**Supp. Fig. 4.**
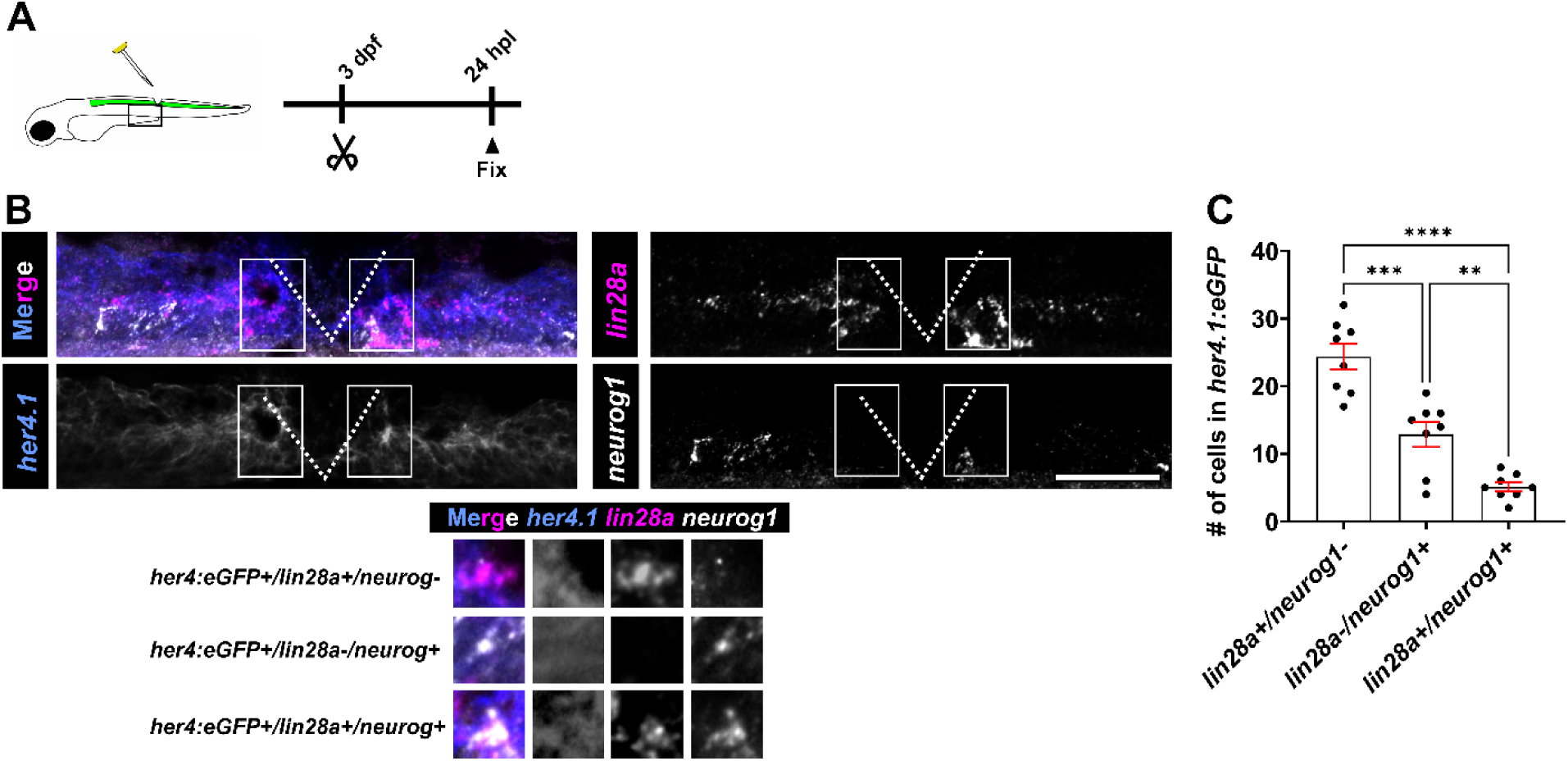
*neurog1* is expressed in a small population of lin28a positive cells. **(A)** Schematic of the experiment preformed in B and quantified in C **(B)** RNAscope against *lin28a* and *neurog1* in *her4.1:eGFP* animals shows sparse triple positive cells. **(C)** Quantification showing that the vast majority of *lin28a^+^* cells are *neurog1^-^* (One-Way ANOVA: p < 0.0001; Tukey’s multiple comparison test: p** = 0.0061, p*** = 0.0001, p**** < 0.0001)**. Scale bars:** 100 µm. Dorsal is up and rostral is left.

**Supp. Fig. 5.**
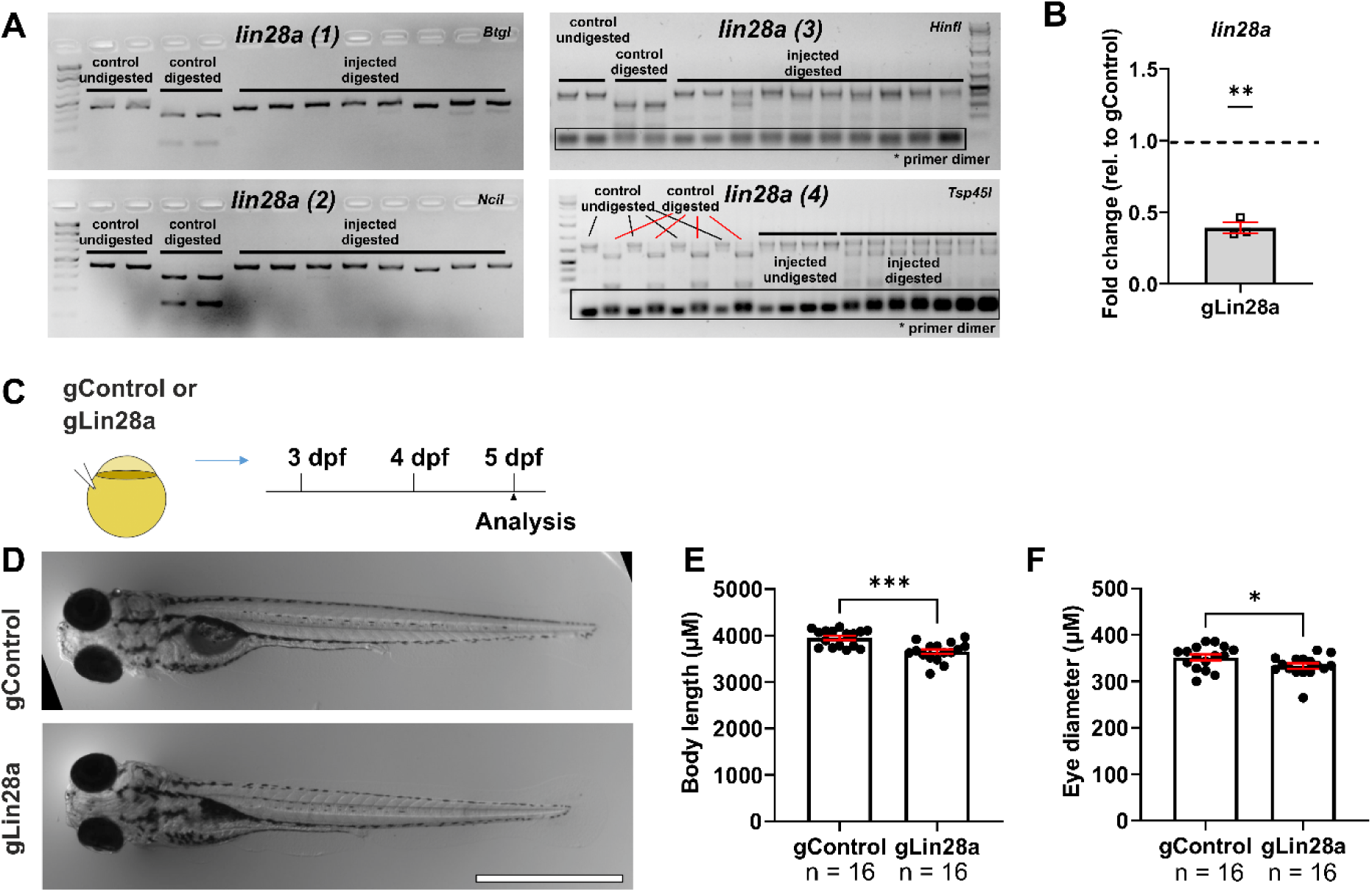
Characterization of *lin28a* somatic mutants. **(A)** RFLP analysis gel for gRNA efficiency validation is shown. Each lane represents one larva with and without digestion with the indicated restriction enzymes and with and without targeting these sites with gRNAs, as indicated. Targeting the recognition sites with haCR gRNAs leads to efficient somatic mutation, as indicated by the almost complete resistance to digestion. **(B)** qPCR showing a reduction of *lin28a* mRNA after gRNA injection**. (C)** A schematic indicating the experimental design for D-F is shown. **(D)** Photomicrographs of gControl and gLin28a larvae are shown at 5 dpf of gRNA1 and 2 injected larvae. **(E-F)** Quantifications show small differences in body length (E: gControl: 3948 µm ± 45.4; gLin28a: 3652 µm ± 51.1; Mann-Whitney U-test: p = 0.0002) and eye diameter in gLin28a larvae, compared tp control gRNA-injecrted animals (F: gControl: 351.6 µm ± 6.3; gLin28a: 333.0 µm ± 5.8; t-test: p = 0.0390). **Scale bars:** 1 mm. Rostral is left.

**Supp. Fig. 6.**
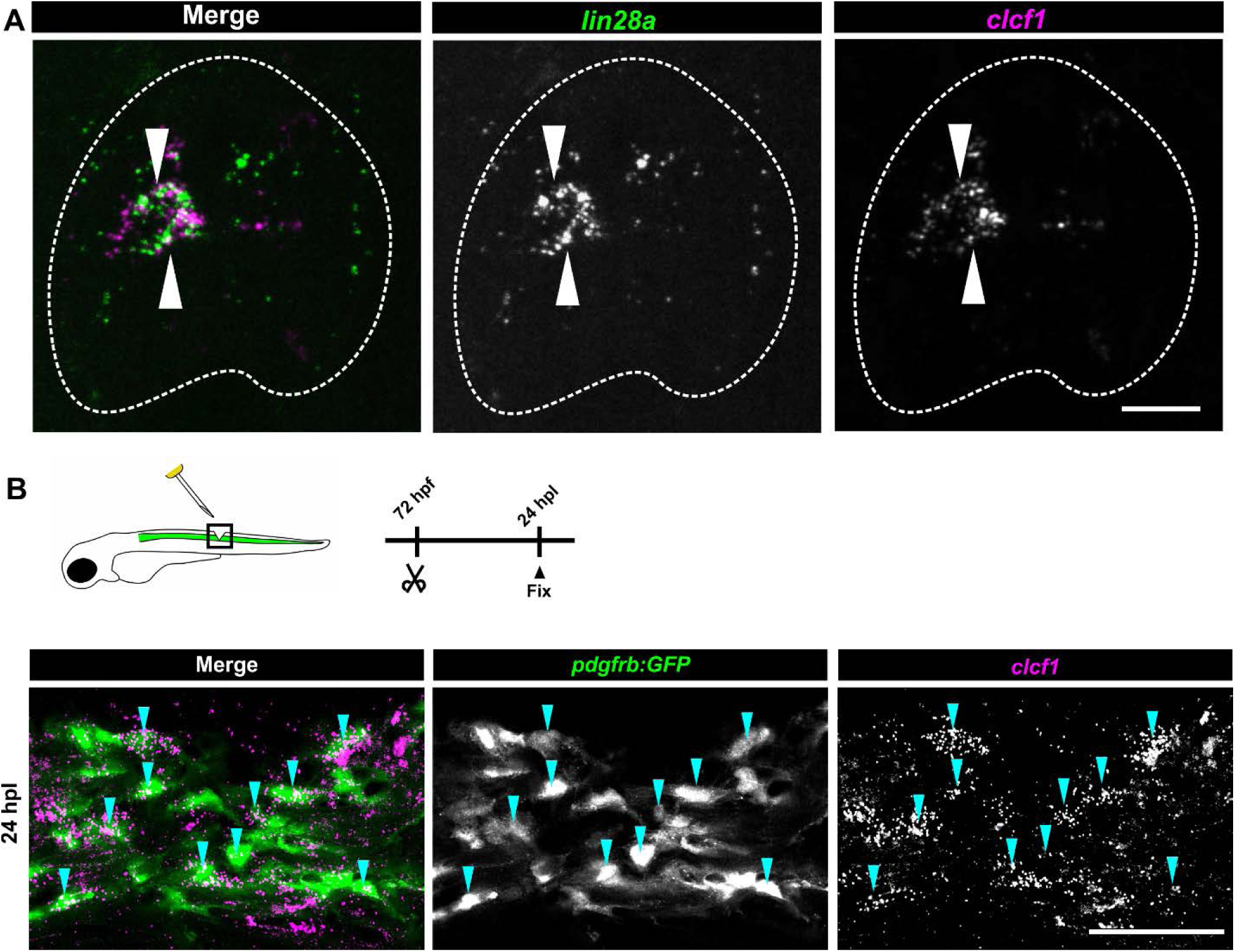
*Clcf1* is co-expressed in *lin28+* cells and also expressed in fibroblasts. **(A)** Cross section of a double HCR-FISH against *lin28a* and *clcf1* showing colocalization of both factors (arrows). **(B)** HCR-FISH against *clcf1* in *pdgfrb:GFP* larvae shows double-positive cells in the injury site (blue arrows). **Scale bar:** 15 µm (A) 100 µm (B).

**Supp. Fig. 7.**
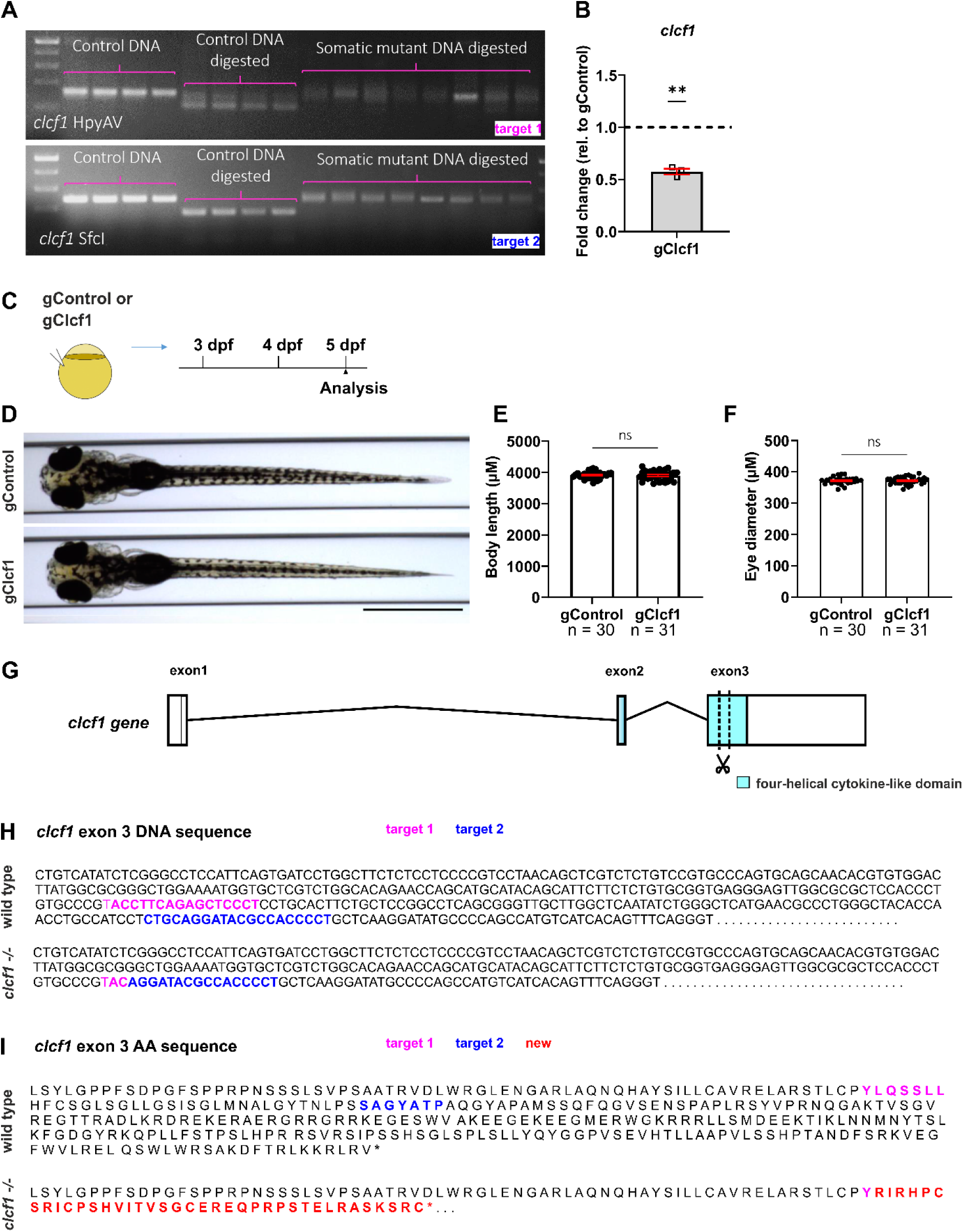
Characterisation of somatic and germline stable *clcf1* mutants. **(A)** RFLP for gRNA efficiency validation of *clcf1* gRNA. Each lane represents one larva with and without digestion with the indicated restriction enzymes and with and without targeting these sites with gRNAs as indicated. Targeting the recognition sites with haCR gRNAs leads to efficient somatic mutation, as indicated by the almost complete resistance to digestion. **(B)** qPCR showing a reduction of *clcf1* mRNA after gRNA injection**. (C)** A schematic indicating the experimental design for D-F is shown. **(D)** Photomicrographs of gControl and gClcf1 larvae are shown at 5 dpf. **(E-F)** Quantifications show no differences in body length (E: gControl: 3917 µm ± 20.7; gClcf1: 3902 µm ± 28.5; t-test: p = 0.6829) or eye diameter in gClcf1 larvae (F: gControl: 372.1 µm ± 2.1; gClcf1: 372.4 µm ± 2.1; t-test: p = 0.9332). **(G)** A schematic of the *clcf1* gene structure showing that the cut area (scissors) in the germline stable mutant disrupts the four helical cytokine-like domain (blue, main identified functional domain of the gene) **(H-I)** DNA and AA sequence of the *clcf1* gene (exon 3) shows the mutation in the *clcf1* germline mutant. Note a large deletion between the two gRNAs injected in F0 that lead to a frame shift (red) and stop codon (asterisk). **Scale bars:** 1 mm. Rostral is left.

**Supp. Fig. 8.**
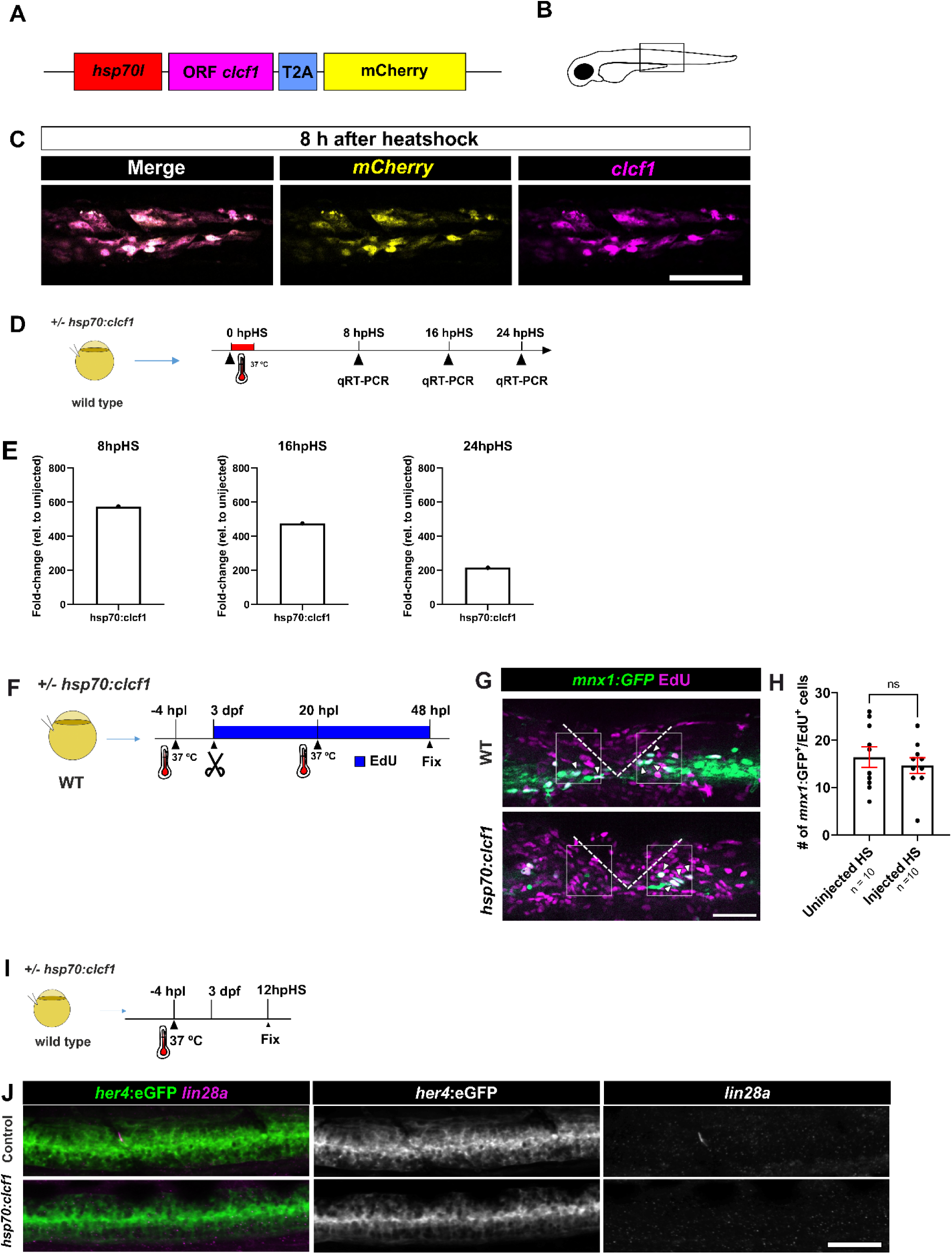
Efficient *clcf1* overexpression has no effect on regenerative neurogenesis. **(A)** Simplified structure of the vector. **(B)** Schematic showing field of view in C. **(C)** HCR-FISH against *mCherry* (yellow) and *clcf1* (magenta) confirms overexpression and colocalization 8h post heat-shock. **(D)** Schematic of experiment performed in E. (**E)** qRT-PCR against *clcf1* at different time points after heat-shock confirms *clcf1* overexpression and a decay in expression over time. **(F)** Schematic of experiment performed in G and quantified in H. **(G)** Over-expression of *clcf1* does not change the number of newly generated neurons after spinal lesion (H; gControl: 16.4 cells per larva ± 2.1; hsp70:clcf1: 14.6 cells per larva ± 1.6; Unpaired t-test: p = 0.5137). **(I-J)** (I) shows schematic of the experiment. (J) HCR-FISH against *lin28a* shows no induced expression of *lin28a* after *clcf1* overexpression in unlesioned larvae. Error bars show SEM. **Scale bars:** 100 µm (C), 50 µm (G, J). Dorsal is up and rostral is left.

**Suppl. Fig. 9.**
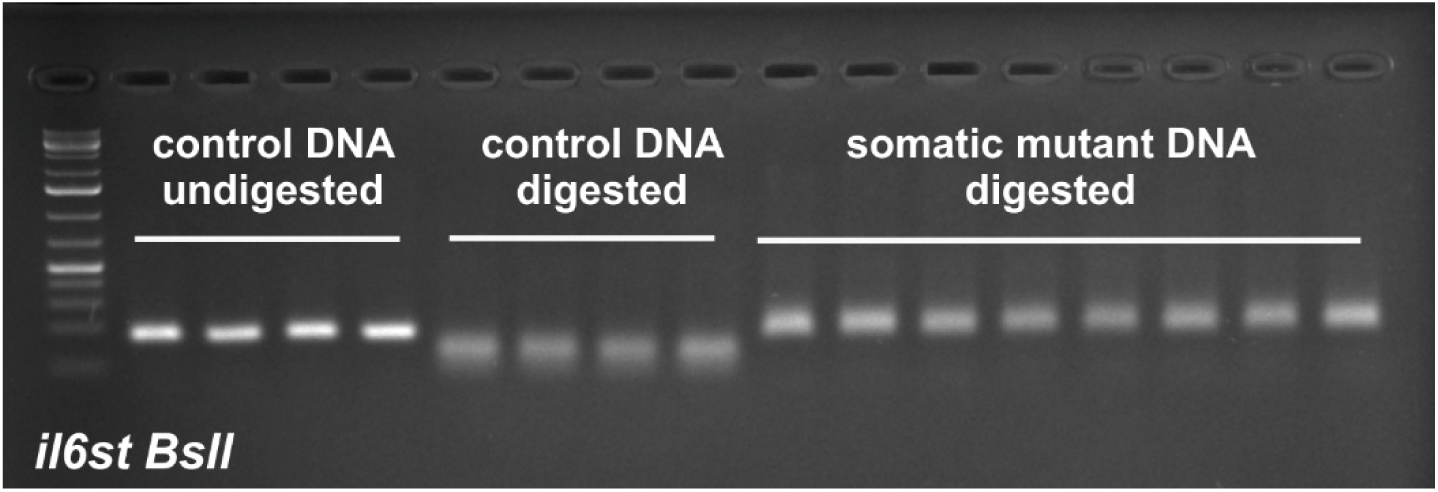
RFLP for *il6st* gRNA efficiency validation. Each lane represents one larva with and without digestion with the indicated restriction enzyme and with and without targeting these sites with gRNAs as indicated. Targeting the recognition sites with haCR gRNAs leads to efficient somatic mutation, as indicated by the almost complete resistance to digestion.

## SUPPLEMENTARY TABLES

**Supplementary Table 1.**
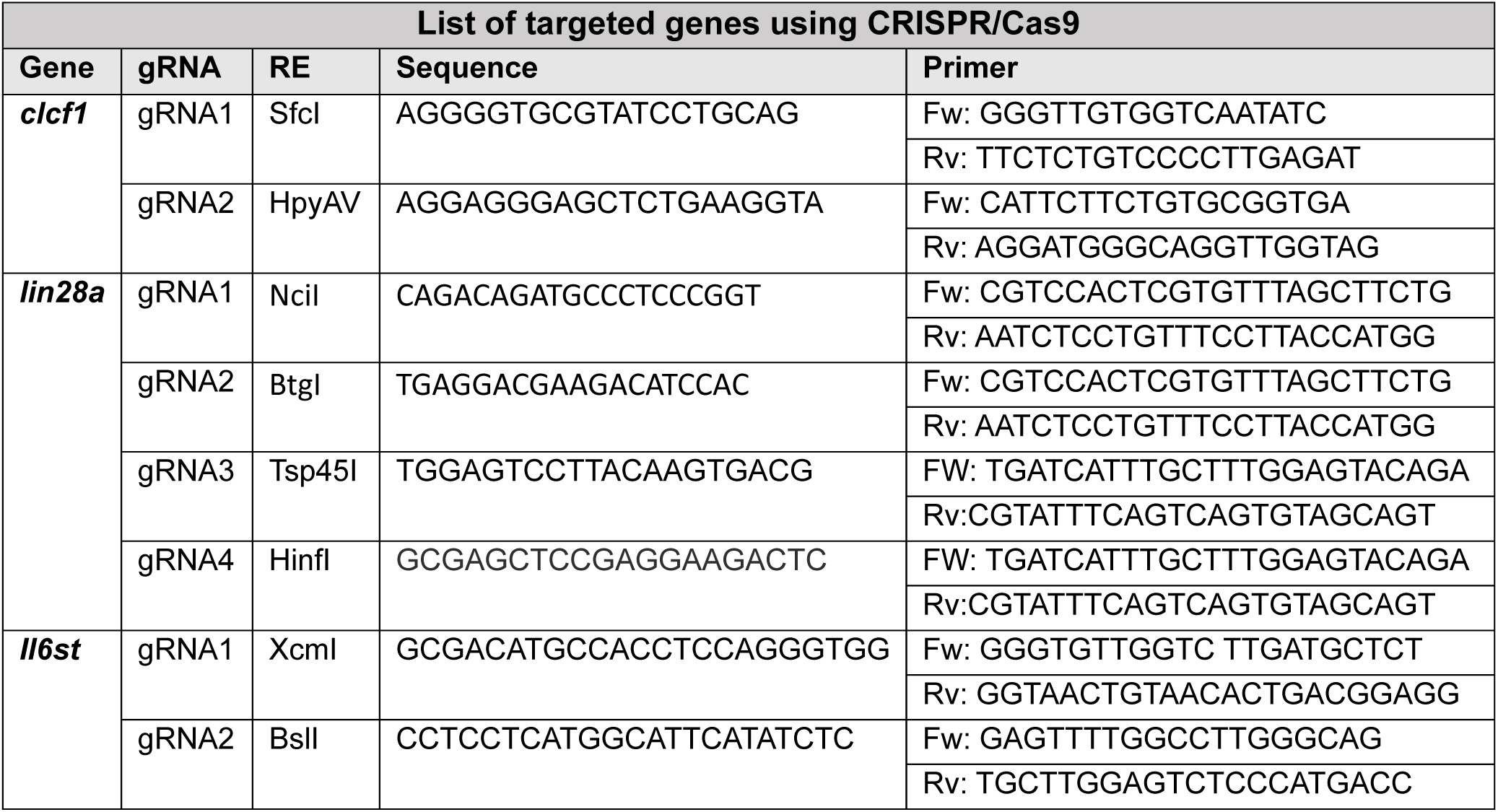
Table showing the gene sequences targeted with gRNAs. Each column represents the targeted sequence of a gene and the primers and restriction enzymes used to validate the efficiency of the mutation.

**Supplementary Table 2.**
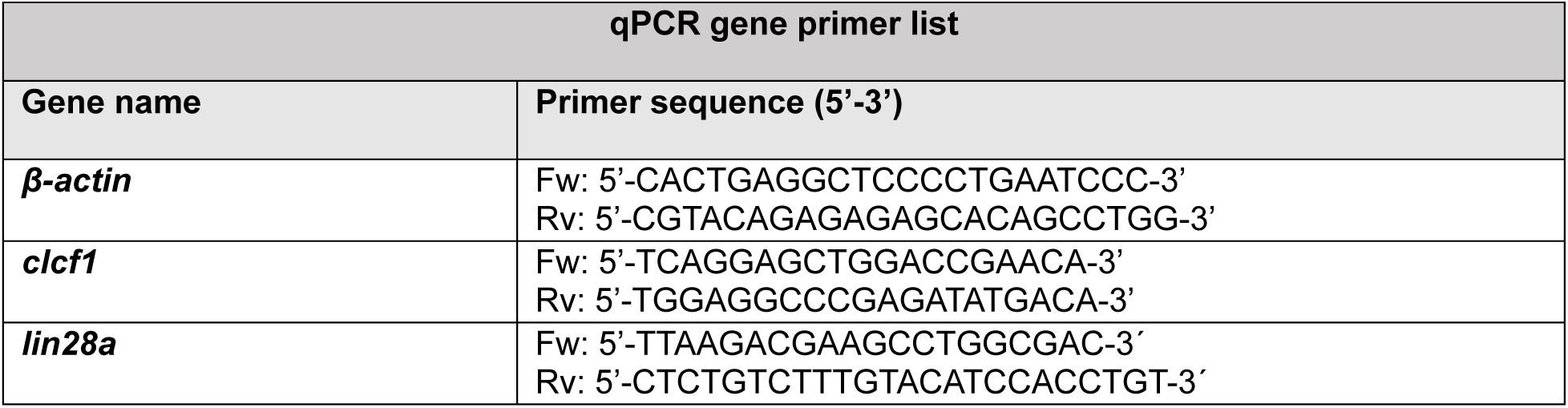
List of qRT-PCR primers.

## REFERENCES

An, Z., Liu, P., Zheng, J., Si, C., Li, T., Chen, Y., Ma, T., Zhang, M. Q., Zhou, Q., & Ding, S. (2019). Sox2 and Klf4 as the Functional Core in Pluripotency Induction without Exogenous Oct4. Cell Rep, 29(7), 1986–2000.e1988. 10.1016/j.celrep.2019.10.026

Ando, K., Fukuhara, S., Izumi, N., Nakajima, H., Fukui, H., Kelsh, R. N., & Mochizuki, N. (2016). Clarification of mural cell coverage of vascular endothelial cells by live imaging of zebrafish. Development, 143(8), 1328–1339. 10.1242/dev.132654

Arber, S., Han, B., Mendelsohn, M., Smith, M., Jessell, T. M., & Sockanathan, S. (1999). Requirement for the homeobox gene Hb9 in the consolidation of motor neuron identity. Neuron, 23(4), 659–674. 10.1016/s0896-6273(01)80026-x

Barnabe-Heider, F., Goritz, C., Sabelstrom, H., Takebayashi, H., Pfrieger, F. W., Meletis, K., & Frisen, J. (2010). Origin of new glial cells in intact and injured adult spinal cord. Cell Stem Cell, 7(4), 470–482. 10.1016/j.stem.2010.07.014

Barreiro-Iglesias, A., Mysiak, K. S., Scott, A. L., Reimer, M. M., Yang, Y., Becker, C. G., & Becker, T. (2015). Serotonin Promotes Development and Regeneration of Spinal Motor Neurons in Zebrafish. Cell Rep, 13(5), 924–932. 10.1016/j.celrep.2015.09.050

Becker, C. G., & Becker, T. (2015). Neuronal regeneration from ependymo-radial glial cells: cook, little pot, cook! Dev Cell, 32(4), 516–527. 10.1016/j.devcel.2015.01.001

Becker, C. G., Lieberoth, B. C., Morellini, F., Feldner, J., Becker, T., & Schachner, M. (2004). L1.1 is involved in spinal cord regeneration in adult zebrafish. J Neurosci, 24(36), 7837–7842. 10.1523/JNEUROSCI.2420-04.2004

Becker, T., Wullimann, M. F., Becker, C. G., Bernhardt, R. R., & Schachner, M. (1997). Axonal regrowth after spinal cord transection in adult zebrafish. J Comp Neurol, 377(4), 577–595. 10.1002/(sici)1096-9861(19970127)377:4<577::aid-cne8>3.0.co;2-#

Bergen, V., Lange, M., Peidli, S., Wolf, F. A., & Theis, F. J. (2020). Generalizing RNA velocity to transient cell states through dynamical modeling. Nat Biotechnol, 38(12), 1408–1414. 10.1038/s41587-020-0591-3

Bernardos, R. L., Barthel, L. K., Meyers, J. R., & Raymond, P. A. (2007). Late-stage neuronal progenitors in the retina are radial Muller glia that function as retinal stem cells. J Neurosci, 27(26), 7028–7040. 10.1523/JNEUROSCI.1624-07.2007

Bertrand, N., Castro, D. S., & Guillemot, F. (2002). Proneural genes and the specification of neural cell types. Nat Rev Neurosci, 3(7), 517–530. 10.1038/nrn874

Bin, G., Jiarong, Z., Shihao, W., Xiuli, S., Cheng, X., Liangbiao, C., & Ming, Z. (2012). Aire promotes the self-renewal of embryonic stem cells through Lin28. Stem Cells Dev, 21(15), 2878–2890. 10.1089/scd.2012.0097

Boyd, P., Campbell, L. J., & Hyde, D. R. (2023). Clcf1/Crlf1a-mediated signaling is neuroprotective and required for Muller glia proliferation in the light-damaged zebrafish retina. Front Cell Dev Biol, 11, 1142586. 10.3389/fcell.2023.1142586

Briona, L. K., Poulain, F. E., Mosimann, C., & Dorsky, R. I. (2015). Wnt/ss-catenin signaling is required for radial glial neurogenesis following spinal cord injury. Dev Biol, 403(1), 15–21. 10.1016/j.ydbio.2015.03.025

Briscoe, J., Pierani, A., Jessell, T. M., & Ericson, J. (2000). A Homeodomain Protein Code Specifies Progenitor Cell Identity and Neuronal Fate in the Ventral Neural Tube. Cell, 101(4), 435–445. 10.1016/S0092-8674(00)80853-3

Casella, G. T., Bunge, M. B., & Wood, P. M. (2006). Endothelial cell loss is not a major cause of neuronal and glial cell death following contusion injury of the spinal cord. Exp Neurol, 202(1), 8–20. 10.1016/j.expneurol.2006.05.028

Castro, D. S., Martynoga, B., Parras, C., Ramesh, V., Pacary, E., Johnston, C., Drechsel, D., Lebel-Potter, M., Garcia, L. G., Hunt, C., Dolle, D., Bithell, A., Ettwiller, L., Buckley, N., & Guillemot, F. (2011). A novel function of the proneural factor Ascl1 in progenitor proliferation identified by genome-wide characterization of its targets. Genes Dev, 25(9), 930–945. 10.1101/gad.627811

Cavone, L., McCann, T., Drake, L. K., Aguzzi, E. A., Oprisoreanu, A. M., Pedersen, E., Sandi, S., Selvarajah, J., Tsarouchas, T. M., Wehner, D., Keatinge, M., Mysiak, K. S., Henderson, B. E. P., Dobie, R., Henderson, N. C., Becker, T., & Becker, C. G. (2021). A unique macrophage subpopulation signals directly to progenitor cells to promote regenerative neurogenesis in the zebrafish spinal cord. Dev Cell, 56(11), 1617–1630 e1616. 10.1016/j.devcel.2021.04.031

Ceto, S., Sekiguchi, K. J., Takashima, Y., Nimmerjahn, A., & Tuszynski, M. H. (2020). Neural Stem Cell Grafts Form Extensive Synaptic Networks that Integrate with Host Circuits after Spinal Cord Injury. Cell stem cell. 10.1016/j.stem.2020.07.007

Charrier, J. B., Lapointe, F., Le Douarin, N. M., & Teillet, M. A. (2002). Dual origin of the floor plate in the avian embryo. Development, 129(20), 4785–4796. 10.1242/dev.129.20.4785

Cigliola, V., Shoffner, A., Lee, N., Ou, J., Gonzalez, T. J., Hoque, J., Becker, C. J., Han, Y., Shen, G., Faw, T. D., Abd-El-Barr, M. M., Varghese, S., Asokan, A., & Poss, K. D. (2023). Spinal cord repair is modulated by the neurogenic factor Hb-egf under direction of a regeneration-associated enhancer. Nat Commun, 14(1), 4857. 10.1038/s41467-023-40486-5

Cimadamore, F., Amador-Arjona, A., Chen, C., Huang, C. T., & Terskikh, A. V. (2013). SOX2-LIN28/let-7 pathway regulates proliferation and neurogenesis in neural precursors. Proc Natl Acad Sci U S A, 110(32), E3017–3026. 10.1073/pnas.1220176110

Cura Costa, E., Otsuki, L., Rodrigo Albors, A., Tanaka, E. M., & Chara, O. (2021). Spatiotemporal control of cell cycle acceleration during axolotl spinal cord regeneration. Elife, 10. 10.7554/eLife.55665

de Winter, J. P., ten Dijke, P., de Vries, C. J., van Achterberg, T. A., Sugino, H., de Waele, P., Huylebroeck, D., Verschueren, K., & van den Eijnden-van Raaij, A. J. (1996). Follistatins neutralize activin bioactivity by inhibition of activin binding to its type II receptors. Mol Cell Endocrinol, 116(1), 105–114. 10.1016/0303-7207(95)03705-5

Dias, T. B., Yang, Y. J., Ogai, K., Becker, T., & Becker, C. G. (2012). Notch signaling controls generation of motor neurons in the lesioned spinal cord of adult zebrafish. J Neurosci, 32(9), 3245–3252. 10.1523/JNEUROSCI.6398-11.2012

Docampo-Seara, A., Cosacak, M. I., Heilemann, K., Kessel, F., Oprişoreanu, A.-M., Westphal, M., Çark, Ö., Zöller, D., Arnold, J., Bretschneider, A., Hnatiuk, A., Ninov, N., Becker, C. G., & Becker, T. (2026). The microglia-derived protein sema4ab attenuates regenerative neurogenesis after spinal cord injury in zebrafish. PLOS Biology, 24(6), e3003865. 10.1371/journal.pbio.3003865

Dusart, I., & Schwab, M. E. (1994). Secondary cell death and the inflammatory reaction after dorsal hemisection of the rat spinal cord. The European journal of neuroscience, 6, 712–724.

Ellis, P., Fagan, B. M., Magness, S. T., Hutton, S., Taranova, O., Hayashi, S., McMahon, A., Rao, M., & Pevny, L. (2004). SOX2, a persistent marker for multipotential neural stem cells derived from embryonic stem cells, the embryo or the adult. Dev Neurosci, 26(2-4), 148–165. 10.1159/000082134

Engel-Pizcueta, C., Hevia, C. F., Voltes, A., Livet, J., & Pujades, C. (2025). Her9 controls the stemness properties of hindbrain boundary cells. Development, 152(1). 10.1242/dev.203164

Ezhkova, D., Schwarzer, S., Spiess, S., Geffarth, M., Machate, A., Zoller, D., Stucke, J., Alexopoulou, D., Lesche, M., Dahl, A., & Hans, S. (2023). Transcriptome analysis reveals an Atoh1b-dependent gene set downstream of Dlx3b/4b during early inner ear development in zebrafish. Biol Open, 12(6). 10.1242/bio.059911

Fausett, B. V., & Goldman, D. (2006). A role for alpha1 tubulin-expressing Muller glia in regeneration of the injured zebrafish retina. J Neurosci, 26(23), 6303–6313. 10.1523/JNEUROSCI.0332-06.2006

Fimbel, S. M., Montgomery, J. E., Burket, C. T., & Hyde, D. R. (2007). Regeneration of inner retinal neurons after intravitreal injection of ouabain in zebrafish. J Neurosci, 27(7), 1712–1724. 10.1523/JNEUROSCI.5317-06.2007

Flanagan-Steet, H., Fox, M. A., Meyer, D., & Sanes, J. R. (2005). Neuromuscular synapses can form in vivo by incorporation of initially aneural postsynaptic specializations. Development, 132(20), 4471–4481. 10.1242/dev.02044

Furtwängler, B., Üresin, N., Richter, S., Schuster, M. B., Barmpouri, D., Holze, H., Wenzel, A., Grønbæk, K., Theilgaard-Mönch, K., Theis, F. J., Schoof, E. M., & Porse, B. T. (2025). Mapping early human blood cell differentiation using single-cell proteomics and transcriptomics. Science, 390(6770), eadr8785. 10.1126/science.adr8785

Goldshmit, Y., Sztal, T. E., Jusuf, P. R., Hall, T. E., Nguyen-Chi, M., & Currie, P. D. (2012). Fgf-dependent glial cell bridges facilitate spinal cord regeneration in zebrafish. J Neurosci, 32(22), 7477–7492. 10.1523/JNEUROSCI.0758-12.2012

Gorsuch, R. A., Lahne, M., Yarka, C. E., Petravick, M. E., Li, J., & Hyde, D. R. (2017). Sox2 regulates Muller glia reprogramming and proliferation in the regenerating zebrafish retina via Lin28 and Ascl1a. Exp Eye Res, 161, 174–192. 10.1016/j.exer.2017.05.012

Gupta, B., Errington, A. C., Jimenez-Pascual, A., Eftychidis, V., Brabletz, S., Stemmler, M. P., Brabletz, T., Petrik, D., & Siebzehnrubl, F. A. (2021). The transcription factor ZEB1 regulates stem cell self-renewal and cell fate in the adult hippocampus. Cell Rep, 36(8), 109588. 10.1016/j.celrep.2021.109588

Hanna, J., Saha, K., Pando, B., van Zon, J., Lengner, C. J., Creyghton, M. P., van Oudenaarden, A., & Jaenisch, R. (2009). Direct cell reprogramming is a stochastic process amenable to acceleration. Nature, 462(7273), 595–601. 10.1038/nature08592

Herrlinger, S., Shao, Q., Yang, M., Chang, Q., Liu, Y., Pan, X., Yin, H., Xie, L.-W., & Chen, J.-F. (2019). Lin28-mediated temporal promotion of protein synthesis is crucial for neural progenitor cell maintenance and brain development in mice. Development, 146(10), dev173765. 10.1242/dev.173765

Johansson, C. B., Momma, S., Clarke, D. L., Risling, M., Lendahl, U., & Frisen, J. (1999). Identification of a neural stem cell in the adult mammalian central nervous system. Cell, 96(1), 25–34. 10.1016/s0092-8674(00)80956-3

Jurisch-Yaksi, N., Yaksi, E., & Kizil, C. (2020). Radial glia in the zebrafish brain: Functional, structural, and physiological comparison with the mammalian glia. Glia, 68(12), 2451–2470. 10.1002/glia.23849

Katreddi, R. R., Taroc, E. Z. M., Hicks, S. M., Lin, J. M., Liu, S., Xiang, M., & Forni, P. E. (2022). Notch signaling determines cell-fate specification of the two main types of vomeronasal neurons of rodents. Development, 149(13). 10.1242/dev.200448

Keatinge, M., Tsarouchas, T. M., Munir, T., Porter, N. J., Larraz, J., Gianni, D., Tsai, H. H., Becker, C. G., Lyons, D. A., & Becker, T. (2021). CRISPR gRNA phenotypic screening in zebrafish reveals pro-regenerative genes in spinal cord injury. PLoS Genet, 17(4), e1009515. 10.1371/journal.pgen.1009515

Knoblich, J. A., Jan, L. Y., & Jan, Y. N. (1995). Asymmetric segregation of Numb and Prospero during cell division. Nature, 377(6550), 624–627. 10.1038/377624a0

Kroll, F., Powell, G. T., Ghosh, M., Gestri, G., Antinucci, P., Hearn, T. J., Tunbak, H., Lim, S., Dennis, H. W., Fernandez, J. M., Whitmore, D., Dreosti, E., Wilson, S. W., Hoffman, E. J., & Rihel, J. (2021). A simple and effective F0 knockout method for rapid screening of behaviour and other complex phenotypes. Elife, 10. 10.7554/eLife.59683

Kuscha, V., Frazer, S. L., Dias, T. B., Hibi, M., Becker, T., & Becker, C. G. (2012a). Lesion-induced generation of interneuron cell types in specific dorsoventral domains in the spinal cord of adult zebrafish. J Comp Neurol, 520(16), 3604–3616. 10.1002/cne.23115

Kuscha, V., Frazer, S. L., Dias, T. B., Hibi, M., Becker, T., & Becker, C. G. (2012b). Lesion-induced generation of interneuron cell types in specific dorsoventral domains in the spinal cord of adult zebrafish. The Journal of comparative neurology, 520(16), 3604–3616. 10.1002/cne.23115

Lee, M. S., Jui, J., Sahu, A., & Goldman, D. (2024). Mycb and Mych stimulate Müller glial cell reprogramming and proliferation in the uninjured and injured zebrafish retina. Development, 151(14). 10.1242/dev.203062

Lee, S. K., Lee, B., Ruiz, E. C., & Pfaff, S. L. (2005). Olig2 and Ngn2 function in opposition to modulate gene expression in motor neuron progenitor cells. Genes Dev, 19(2), 282–294. 10.1101/gad.1257105

Li, Y., Li, A., Glas, M., Lal, B., Ying, M., Sang, Y., Xia, S., Trageser, D., Guerrero-Cázares, H., Eberhart, C. G., Quiñones-Hinojosa, A., Scheffler, B., & Laterra, J. (2011). c-Met signaling induces a reprogramming network and supports the glioblastoma stem-like phenotype. Proc Natl Acad Sci U S A, 108(24), 9951–9956. 10.1073/pnas.1016912108

Llorens-Bobadilla, E., Chell, J. M., Le Merre, P., Wu, Y., Zamboni, M., Bergenstrahle, J., Stenudd, M., Sopova, E., Lundeberg, J., Shupliakov, O., Carlen, M., & Frisen, J. (2020). A latent lineage potential in resident neural stem cells enables spinal cord repair. Science, 370(6512). 10.1126/science.abb8795

Lu, Q. R., Sun, T., Zhu, Z., Ma, N., Garcia, M., Stiles, C. D., & Rowitch, D. H. (2002). Common developmental requirement for Olig function indicates a motor neuron/oligodendrocyte connection. Cell, 109(1), 75–86. 10.1016/s0092-8674(02)00678-5

Luo, Z., Gao, X., Lin, C., Smith, E. R., Marshall, S. A., Swanson, S. K., Florens, L., Washburn, M. P., & Shilatifard, A. (2015). Zic2 is an enhancer-binding factor required for embryonic stem cell specification. Mol Cell, 57(4), 685–694. 10.1016/j.molcel.2015.01.007

Lust, K., Maynard, A., Gomes, T., Fleck, J. S., Camp, J. G., Tanaka, E. M., & Treutlein, B. (2022). Single-cell analyses of axolotl telencephalon organization, neurogenesis, and regeneration. Science, 377(6610), eabp9262. 10.1126/science.abp9262

Meletis, K., Barnabe-Heider, F., Carlen, M., Evergren, E., Tomilin, N., Shupliakov, O., & Frisen, J. (2008). Spinal cord injury reveals multilineage differentiation of ependymal cells. PLoS Biol, 6(7), e182. 10.1371/journal.pbio.0060182

Mitra, S., Sharma, P., Kaur, S., Khursheed, M. A., Gupta, S., Chaudhary, M., Kurup, A. J., & Ramachandran, R. (2019). Dual regulation of lin28a by Myc is necessary during zebrafish retina regeneration. J Cell Biol, 218(2), 489–507. 10.1083/jcb.201802113

Mokalled, M. H., Patra, C., Dickson, A. L., Endo, T., Stainier, D. Y., & Poss, K. D. (2016). Injury-induced ctgfa directs glial bridging and spinal cord regeneration in zebrafish. Science, 354(6312), 630–634. 10.1126/science.aaf2679

Moldovan, G. L., Pfander, B., & Jentsch, S. (2007). PCNA, the maestro of the replication fork. Cell, 129(4), 665–679. 10.1016/j.cell.2007.05.003

Morris, R., Kershaw, N. J., & Babon, J. J. (2018). The molecular details of cytokine signaling via the JAK/STAT pathway. Protein Sci, 27(12), 1984–2009. 10.1002/pro.3519

Moss, E. G., Lee, R. C., & Ambros, V. (1997). The cold shock domain protein LIN-28 controls developmental timing in C. elegans and is regulated by the lin-4 RNA. Cell, 88(5), 637–646. 10.1016/s0092-8674(00)81906-6

Moss, E. G., & Tang, L. (2003). Conservation of the heterochronic regulator Lin-28, its developmental expression and microRNA complementary sites. Dev Biol, 258(2), 432–442. 10.1016/s0012-1606(03)00126-x

Nüsslein-Volhard, C., & Dahm, R. (2002). Zebrafish: a practical approach. Oxford University Press.

Ohnmacht, J., Yang, Y., Maurer, G. W., Barreiro-Iglesias, A., Tsarouchas, T. M., Wehner, D., Sieger, D., Becker, C. G., & Becker, T. (2016). Spinal motor neurons are regenerated after mechanical lesion and genetic ablation in larval zebrafish. Development, 143(9), 1464–1474. 10.1242/dev.129155

Ouchi, Y., Yamamoto, J., & Iwamoto, T. (2014). The heterochronic genes lin-28a and lin-28b play an essential and evolutionarily conserved role in early zebrafish development. PLoS One, 9(2), e88086. 10.1371/journal.pone.0088086

Park, H. C., Shin, J., & Appel, B. (2004). Spatial and temporal regulation of ventral spinal cord precursor specification by Hedgehog signaling. Development, 131(23), 5959–5969. 10.1242/dev.01456

Pecce, V., Verrienti, A., Fiscon, G., Sponziello, M., Conte, F., Abballe, L., Durante, C., Farina, L., Filetti, S., & Paci, P. (2021). The role of FOSL1 in stem-like cell reprogramming processes. Sci Rep, 11(1), 14677. 10.1038/s41598-021-94072-0

Ramachandran, R., Fausett, B. V., & Goldman, D. (2010). Ascl1a regulates Muller glia dedifferentiation and retinal regeneration through a Lin-28-dependent, let-7 microRNA signalling pathway. Nat Cell Biol, 12(11), 1101–1107. 10.1038/ncb2115

Raymond, P. A., Barthel, L. K., Bernardos, R. L., & Perkowski, J. J. (2006). Molecular characterization of retinal stem cells and their niches in adult zebrafish. BMC Dev Biol, 6, 36. 10.1186/1471-213X-6-36

Reimer, M. M., Kuscha, V., Wyatt, C., Sorensen, I., Frank, R. E., Knuwer, M., Becker, T., & Becker, C. G. (2009). Sonic hedgehog is a polarized signal for motor neuron regeneration in adult zebrafish. J Neurosci, 29(48), 15073–15082. 10.1523/jneurosci.4748-09.2009

Reimer, M. M., Norris, A., Ohnmacht, J., Patani, R., Zhong, Z., Dias, T. B., Kuscha, V., Scott, A. L., Chen, Y. C., Rozov, S., Frazer, S. L., Wyatt, C., Higashijima, S., Patton, E. E., Panula, P., Chandran, S., Becker, T., & Becker, C. G. (2013). Dopamine from the brain promotes spinal motor neuron generation during development and adult regeneration. Dev Cell, 25(5), 478–491. 10.1016/j.devcel.2013.04.012

Reimer, M. M., Sorensen, I., Kuscha, V., Frank, R. E., Liu, C., Becker, C. G., & Becker, T. (2008). Motor neuron regeneration in adult zebrafish. J Neurosci, 28(34), 8510–8516. 10.1523/JNEUROSCI.1189-08.2008

Ribeiro, A., Monteiro, J. F., Certal, A. C., Cristovão, A. M., & Saúde, L. (2017). Foxj1a is expressed in ependymal precursors, controls central canal position and is activated in new ependymal cells during regeneration in zebrafish. Open Biol, 7(11). 10.1098/rsob.170139

Rodrigo Albors, A., Tazaki, A., Rost, F., Nowoshilow, S., Chara, O., & Tanaka, E. M. (2015). Planar cell polarity-mediated induction of neural stem cell expansion during axolotl spinal cord regeneration. Elife, 4, e10230. 10.7554/eLife.10230

Rondon, R., Hezez, T., Richard Albert, J., Hayashi, S., Drayton-Libotte, B., Curto, G. G., Auradé, F., Balloul, E., Dugast-Darzacq, C., Relaix, F., Gilardi-Hebenstreit, P., & Ribes, V. (2025). Dual transcriptional activities of PAX3 and PAX7 spatially encode spinal cell fates through distinct gene networks. PLoS Biol, 23(10), e3003448. 10.1371/journal.pbio.3003448

Rose-John, S. (2018). Interleukin-6 Family Cytokines. Cold Spring Harb Perspect Biol, 10(2). 10.1101/cshperspect.a028415

Ross, S. E., Greenberg, M. E., & Stiles, C. D. (2003). Basic helix-loop-helix factors in cortical development. Neuron, 39(1), 13–25. 10.1016/s0896-6273(03)00365-9

Rost, F., Rodrigo Albors, A., Mazurov, V., Brusch, L., Deutsch, A., Tanaka, E. M., & Chara, O. (2016). Accelerated cell divisions drive the outgrowth of the regenerating spinal cord in axolotls. Elife, 5. 10.7554/eLife.20357

Rueden, C. T., Schindelin, J., Hiner, M. C., DeZonia, B. E., Walter, A. E., Arena, E. T., & Eliceiri, K. W. (2017). ImageJ2: ImageJ for the next generation of scientific image data. BMC Bioinformatics, 18(1), 529. 10.1186/s12859-017-1934-z

Sabelstrom, H., Stenudd, M., & Frisen, J. (2014). Neural stem cells in the adult spinal cord. Exp Neurol, 260, 44–49. 10.1016/j.expneurol.2013.01.026

Scardigli, R., Schuurmans, C., Gradwohl, G., & Guillemot, F. (2001). Crossregulation between Neurogenin2 and pathways specifying neuronal identity in the spinal cord. Neuron, 31(2), 203–217. 10.1016/s0896-6273(01)00358-0

Schindelin, J., Arganda-Carreras, I., Frise, E., Kaynig, V., Longair, M., Pietzsch, T., Preibisch, S., Rueden, C., Saalfeld, S., Schmid, B., Tinevez, J. Y., White, D. J., Hartenstein, V., Eliceiri, K., Tomancak, P., & Cardona, A. (2012). Fiji: an open-source platform for biological-image analysis. Nat Methods, 9(7), 676-682. 10.1038/nmeth.2019

Scholzen, T., & Gerdes, J. (2000). The Ki-67 protein: from the known and the unknown. J Cell Physiol, 182(3), 311–322. 10.1002/(sici)1097-4652(200003)182:3<311::Aid-jcp1>3.0.Co;2-9

Sharma, P., Gupta, S., Chaudhary, M., Mitra, S., Chawla, B., Khursheed, M. A., & Ramachandran, R. (2019). Oct4 mediates Muller glia reprogramming and cell cycle exit during retina regeneration in zebrafish. Life Sci Alliance, 2(5). 10.26508/lsa.201900548

Shin, J., Park, H. C., Topczewska, J. M., Mawdsley, D. J., & Appel, B. (2003). Neural cell fate analysis in zebrafish using olig2 BAC transgenics. Methods Cell Sci, 25(1-2), 7–14. http://www.ncbi.nlm.nih.gov/entrez/query.fcgi?cmd=Retrieve&db=PubMed&dopt=Citation&list_uids=14739582

Silver, J., & Miller, J. H. (2004). Regeneration beyond the glial scar. Nat Rev Neurosci, 5(2), 146–156. 10.1038/nrn1326

Velasco, S., Ibrahim, M. M., Kakumanu, A., Garipler, G., Aydin, B., Al-Sayegh, M. A., Hirsekorn, A., Abdul-Rahman, F., Satija, R., Ohler, U., Mahony, S., & Mazzoni, E. O. (2017). A Multi-step Transcriptional and Chromatin State Cascade Underlies Motor Neuron Programming from Embryonic Stem Cells. Cell Stem Cell, 20(2), 205–217 e208. 10.1016/j.stem.2016.11.006

Walder, S., Zhang, F., & Ferretti, P. (2003). Up-regulation of neural stem cell markers suggests the occurrence of dedifferentiation in regenerating spinal cord. Dev Genes Evol, 213(12), 625–630. 10.1007/s00427-003-0364-2

Weiss, S., Dunne, C., Hewson, J., Wohl, C., Wheatley, M., Peterson, A. C., & Reynolds, B. A. (1996). Multipotent CNS stem cells are present in the adult mammalian spinal cord and ventricular neuroaxis. J Neurosci, 16(23), 7599–7609. 10.1523/JNEUROSCI.16-23-07599.1996

Westerfield, M. (2000). *The zebrafish book: a guide for the laboratory use of zebrafish (Danio rerio)* (4th ed.). University of Oregon Press.

Yang, D. H., & Moss, E. G. (2003). Temporally regulated expression of Lin-28 in diverse tissues of the developing mouse. Gene Expr Patterns, 3(6), 719–726. 10.1016/s1567-133x(03)00140-6

Yang, M., Yang, S. L., Herrlinger, S., Liang, C., Dzieciatkowska, M., Hansen, K. C., Desai, R., Nagy, A., Niswander, L., Moss, E. G., & Chen, J. F. (2015). Lin28 promotes the proliferative capacity of neural progenitor cells in brain development. Development, 142(9), 1616–1627. 10.1242/dev.120543

Yeo, S. Y., Kim, M., Kim, H. S., Huh, T. L., & Chitnis, A. B. (2007). Fluorescent protein expression driven by her4 regulatory elements reveals the spatiotemporal pattern of Notch signaling in the nervous system of zebrafish embryos. Dev Biol, 301(2), 555–567. 10.1016/j.ydbio.2006.10.020

Yu, J., Vodyanik, M. A., Smuga-Otto, K., Antosiewicz-Bourget, J., Frane, J. L., Tian, S., Nie, J., Jonsdottir, G. A., Ruotti, V., Stewart, R., Slukvin, II, & Thomson, J. A. (2007). Induced pluripotent stem cell lines derived from human somatic cells. Science, 318(5858), 1917–1920. 10.1126/science.1151526

Zhao, X. F., Wan, J., Powell, C., Ramachandran, R., Myers, M. G., Jr., & Goldman, D. (2014). Leptin and IL-6 family cytokines synergize to stimulate Muller glia reprogramming and retina regeneration. Cell Rep, 9(1), 272–284. 10.1016/j.celrep.2014.08.047

